# Comparative analysis of epigenetic aging clocks from CpG characteristics to functional associations

**DOI:** 10.1101/512483

**Authors:** Zuyun Liu, Diana Leung, Morgan Levine

**Author notes:** **Correspondence:** Morgan Levine, Ph.D., Department of Pathology, 300 George Street, Yale School of Medicine, New Haven, CT 06511, USA.

## Abstract

To date, a number of epigenetic clocks have been developed using DNA methylation data, aimed at approximating biological aging in multiple tissues/cells. However, despite the assumption that these clocks are meant to capture the same phenomenon-aging, their correlations with each other are weak, and there is a lack of consistency in their associations with outcomes of aging. Therefore, the goal of this study was to compare and contrast the molecular characteristics and functional associations of 11 existing epigenetic clocks, using data from diverse human tissue and cell types. Results suggest that the CpGs comprised in the various clocks differ in regards to the consistency of their age correlations across tissues/cells. Using microarray expression data from purified CD14+ monocytes, we found that six clocks—Yang, Hannum, Lin, Levine, Horvath1, and Horvath2—has relatively similar transcriptional profiles. Network analysis revealed nine co-expression modules, most of which display robust correlations across various clocks. One significant module—turquoise is involved in mitochondrial translation, gene expression, respiratory chain complex assembly, and oxidative phosphorylation. Finally, using data from 143B cells with chronically depleted mtDNA (rho0) and 143B controls, we found that rho0 cells have more than a three-standard deviation increase in epigenetic age for Levine (p=0.006), Lin (p=0.012), and Yang (p=0.013). In summary, these results demonstrate the shared and contrasting features of existing epigenetic clocks, in regards to the CpG characteristic, tissue specificity, and co-regulatory gene network signatures, and suggesting a link between two hallmarks of aging—epigenetic alterations and mitochondrial dysfunction.

## Introduction

Chronological age is arguably the strongest risk factor for most major chronic diseases [1], suggesting may be a causal driver of pathogenesis for various diseases [2]. In response, there has been a major initiative focused on targeting aging directly in order to increase disease-free life expectancy and in turn reduce healthcare spending and improve population health. However, given that aging is a complex multifactorial process characterized by increasing dysregulation and loss of function across multiple levels and systems [3], one major hurdle in tackling human aging is how to measure it. Thus, quantify aging, particularly using molecular hallmarks, has become a priority in Geroscience research [2, 4, 5].

Measures based on epigenetic alterations have emerged as promising biomarkers of aging, in part stemming from the accumulating evidence that the methylome exhibits extremely precise transformations with age [6-9]. Such changes have also been identified as potential therapeutic targets, in line with the Geroscience paradigm proposed by the National Institute on Aging [2]. In particular, DNA methylation (DNAm)—typically consisting of the covalent attachment of methyl groups to cytosines in CpG dinucleotides—has exhibited extremely robust age tends, characterized by specific CpG island (CGI) promotor-associated hypermethylation, and genome-wide hypomethylation. Building off the precision of age-related DNAm changes, a number of methylation-based age predictors, commonly referred to as epigenetic clocks or DNAm age, have been developed (a brief review of these epigenetic clocks can be found in Additional file 1 and other literature [10]). These epigenetic clocks, which exhibit extremely high correlations with chronological age, upwards of r=0.95 in full age range samples, are thought to capture aspects of biological aging, or more accurately epigenetic aging. Further, these epigenetic clocks can be contrasted against individuals’ chronological ages to capture interindividual and/or inter-tissue variability in the rate of aging [11]. For instance, samples predicted to have a higher epigenetic age, relative to chronological age, are characterized as exhibiting signs of accelerated aging. For a number of the clocks, this divergence between epigenetic and chronological age has been shown to translate to differential susceptibility to death and disease. After adjustment for chronological age, epigenetic age often remains a very significant predictor of mortality [12-15]. Additionally, some clocks have been shown to predict major diseases of aging, including coronary heart disease, diabetes, and some forms of cancer (e.g., breast and lung) [15-19]. Moreover, differential epigenetic age in brain is associated with both Alzheimer’s related neuropathology and Down syndrome. Finally, there is also evidence to suggest some clocks reflect HIV infection, obesity, Hutchinson Gilford Progeria Syndrome [20], and familial longevity [15, 21].

However, despite the general associations between epigenetic clocks and outcomes of aging, the strengths of these associations vary significantly across different clocks. Moreover, after adjusting for chronological age, the majority of the clocks are typically only correlated at r<0.5—suggesting that they may be capturing different aspects of epigenetic aging. Furthermore, given that each clock utilized diverse sample types when being developed, their translatability in other tissues/cells or at various developmental stages differs. As a result, the goal of this study was to compare and contrast the CpG characteristics, tissue-specificity, and transcriptional signals of 11 published epigenetic clocks, in order to identify their shared and/or distinct biological etiology.

## Results

### Clocks vary in the types of CpGs they contain

The epigenetic clocks we considered are listed in Table 1. For clocks comparisons, we started by describing CpG characteristics (Fig 1). Overall, about 1,600 CpGs are included when pooling across all clocks. The majority of CpGs (n=1,427) are unique to only one clock and show no overlap. Conversely, one CpG (cg09809672) overlaps in seven clocks, and 37 CpGs overlap in three to five clocks. Moreover, this does not change when considering larger genomic regions, suggesting that clocks are not simply selecting adjacent/collinear CpGs and instead are drawing markers from entirely different genomic regions. This difference is further emphasized when comparing CpG annotations (see Fig S1A in Additional file 2 and 3). Four clocks (Hannum, Horvath1, Lin, and Levine) show relatively similar proportions in regards to CpG locations, such that islands, shores, and shelves/open seas each made up about a third of these clocks CpGs. The Horvath2 clock is somewhat similar, although with slightly fewer CpGs in islands, and slightly more in Open Seas. Conversely, the Yang clock is comprised almost exclusively of CpGs in islands (85%), while the Zhang clock and the Vidal-Bralo clock contain only one CpG located in CGIs.

**Table 1.**
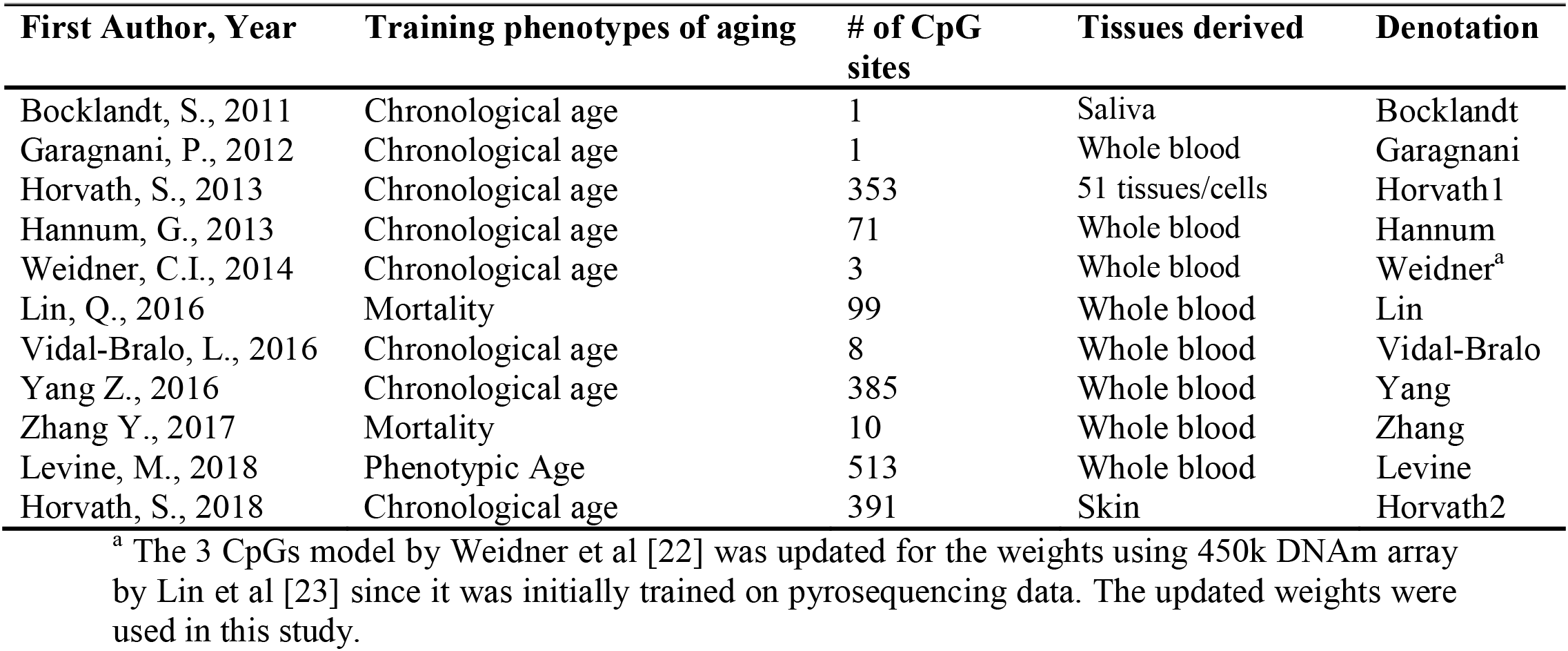
Summary of recent existing 11 epigenetic aging clocks in human being

| First Author, Year | Training phenotypes of aging | # of CpG sites | Tissues derived | Denotation |
| --- | --- | --- | --- | --- |
| Bocklandt, S., 2011 | Chronological age | 1 | Saliva | Bocklandt |
| Garagnani, P., 2012 | Chronological age | 1 | Whole blood | Garagnani |
| Horvath, S., 2013 | Chronological age | 353 | 51 tissues/cells | Horvath1 |
| Hannum, G., 2013 | Chronological age | 71 | Whole blood | Hannum |
| Weidner, C.I., 2014 | Chronological age | 3 | Whole blood | Weidner<sup>a</sup> |
| Lin, Q., 2016 | Mortality | 99 | Whole blood | Lin |
| Vidal-Bralo, L., 2016 | Chronological age | 8 | Whole blood | Vidal-Bralo |
| Yang Z., 2016 | Chronological age | 385 | Whole blood | Yang |
| Zhang Y., 2017 | Mortality | 10 | Whole blood | Zhang |
| Levine, M., 2018 | Phenotypic Age | 513 | Whole blood | Levine |
| Horvath, S., 2018 | Chronological age | 391 | Skin | Horvath2 |

<sup>a</sup> The 3 CpGs model by Weidner et al [22] was updated for the weights using 450k DNAm array by Lin et al [23] since it was initially trained on pyrosequencing data. The updated weights were used in this study.

**Fig 1.**
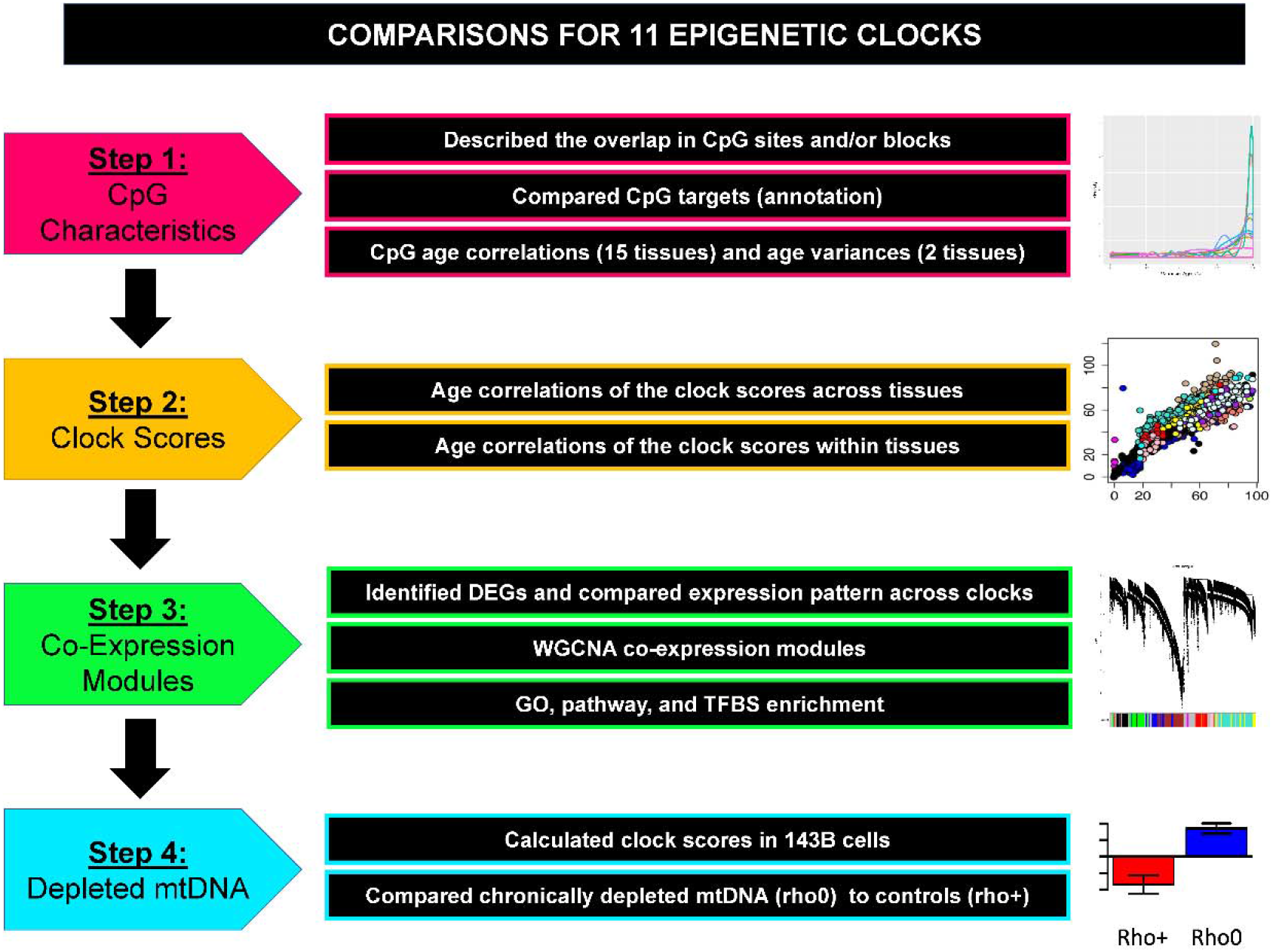
The analytic plan for this study. DEG, differentially expressed genes; WGCNA, weighted-gene correlation network analysis; GO, Gene Ontology; TFBS, transcription factor binding site.

These clocks also vary in regards to the proportions of CpGs located in regions marked by polycomb-group (PcG) protein targets and DNase I hypersensitive sites (DHS) (Fig S1B and S1C), implying the CpGs contained in them may have differing regulatory mechanisms. For instance, Yang includes only CpGs in PcG protein targets, whereas, for the most part, CpGs in PcG protein targets make less than a third of CpGs in the other clocks (range 12-30%). Similarly, the Yang clock also has the highest proportion of DHS CpGs (42%), followed by Hannum with 37%, Zhang with 30%, Horvath2 with 25%, Lin with 18%, Levine and Horvath1 with about 13%, and both Vidal-Bralo and Weidner containing none.

### CpGs track age across tissues and exhibit increasing noise with age

Next, we examined the strength and tissue-specificity of the age correlations for the individual CpGs in each clock (Fig 2). We observed that most of the CpGs in the Hannum clock show strong and consistent age correlation regardless of tissue and/or cell type, with clear distinctions between approximately half of the CpGs showing positive age correlations and the other half showing negative age correlations. Lin, Horvath2, and Vidal-Bralo exhibit similar tissue consistency, but with slightly weaker correlations and also few positively age-associated CpGs. Horvath1 and Levine exhibit good tissue consistency, but contain a large proportion of CpGs with very weak to no age correlation. Yang almost exclusively contains CpGs with positive age correlations; however, the strength tends to be somewhat weaker than what is observed for Hannum, Horvath2, or Lin. Lastly, we observed high tissue specificity for the CpGs in the Zhang clock.

**Fig 2.**
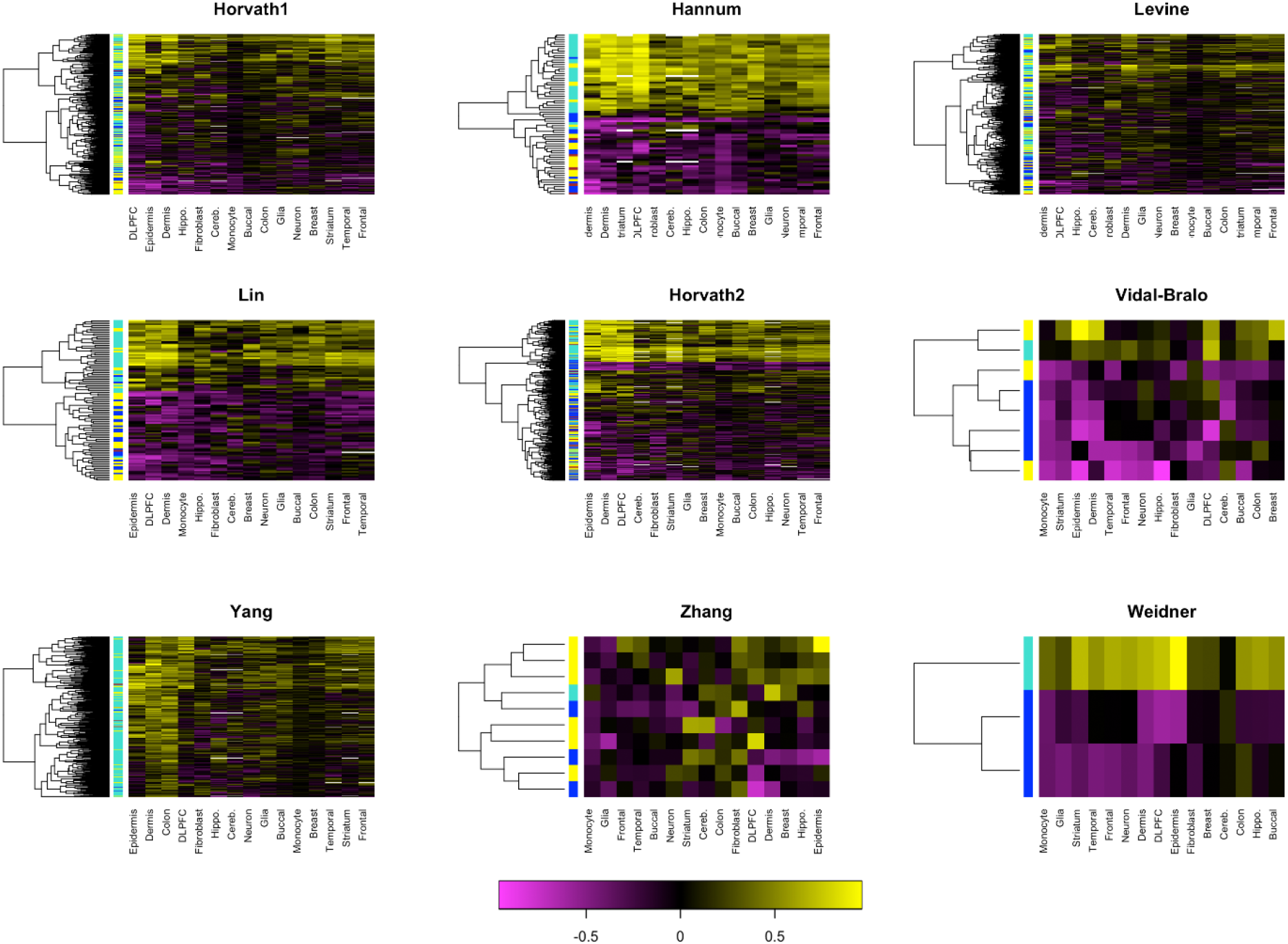
Heatmaps (hierarchical clustering) of age correlations for the CpGs included in each epigenetic clock, across various tissues and cells. DLPFX, dorsolateral prefrontal cortex.

We also examined the change in variance with age for each CpG, which can be thought of as increasing noise or heterogeneity in DNAm across cells. Changes in DNAm that represent random drift should exhibit increasing variance with age; conversely, DNAm changes that are more developmental should exhibit stable variance over the age range. Using data from purified monocytes and DLPFC, we estimated the correlations between the variance in CpG DNAm levels within 10-year age bins (e.g., ages 30-39, 40-49) and the midpoint for age in each bin. We found that Lin and Hannum almost exclusively contain CpGs that display strong increases in variance of DNAm in monocytes with age (Fig 3A). The other clocks, aside from Zhang, mainly consist of CpGs for which variance increases with age in monocytes; however, to a lesser degree than is seen for Lin and Hannum. When examining changes in variance using data from DLPFC, the age trend is substantially reduced (Fig 3B). In this case the Weidner (3 CpGs) and Vidal-Bralo (8 CpGs) clocks show general consensus for increasing variance with age, while the other clocks have only a slight enrichment in CpGs that exhibit increasing DNAm variance with age.

**Fig 3.**
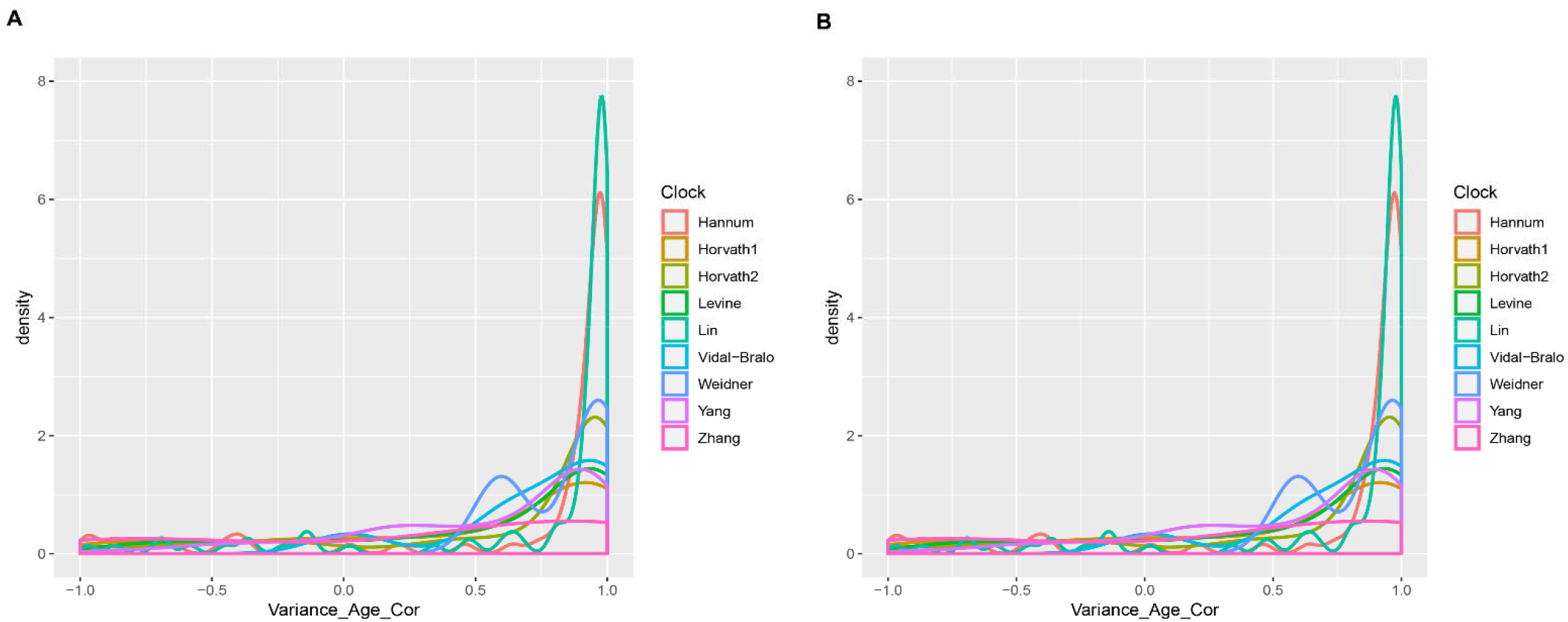
Variance of each CpG using 10-year age bins in A. purified monocytes and B. DLPFC datasets. DLPFX, dorsolateral prefrontal cortex. Positive values suggest that variance increases with age, whereas negative values suggest variance decreases with age.

### Epigenetic clocks track age across tissues

Fig 4 illustrates the age correlations for the clock scores across tissue and cell types. As expected the original pan-tissue clock by Horvath (Horvath1) has the strongest age correlation across pooled tissues and cells (r=0.94). This is followed by Horvath2 (r=0.85), which was the only other age predictor trained on more than one tissue type. Nevertheless, despite being trained in whole blood, the clocks by Hannum, Levine, Lin, and Weidner also exhibit fairly robust multi-tissue age correlations (r=0.68, 0.53, 0.67, and 0.46, respectively). Further, the loss in the strength of the correlations appear to be due to tissue differences rather than a lack of age association within tissues, as suggested by Fig S2 (Additional file 4), which shows that the within tissue age correlations are extremely strong for these clocks. Furthermore, age slopes in these clocks appear to be differentiated by brain versus non-brain samples—with brain showing a much slower increase in epigenetic age over chronological age. Interestingly, the single CpG clock by Garagnani exhibits similarly robust age correlations both across tissues (r=0.70) and within tissues. Much weaker multi-tissue age correlations were observed for the Vidal-Bralo clock (r=0.11) and the Yang clock (r=-0.23). However, the Yang clock tends to exhibit a very strong positive age slope in colon (yellow) and epidermis (pink), which is not surprising given that Yang was developed to be a mitotic clock. As with the other clocks, Yang and Vidal-Bralo reflect tissue differences, but show moderately robust correlations within tissues. The single CpG clock by Bocklandt exhibits stark tissue differences, but moderate within tissue correlations. Finally, the clock by Zhang does not show good consensus regarding its age correlation across tissues, and in fact is negative in some tissues and positive in others.

**Fig 4.**
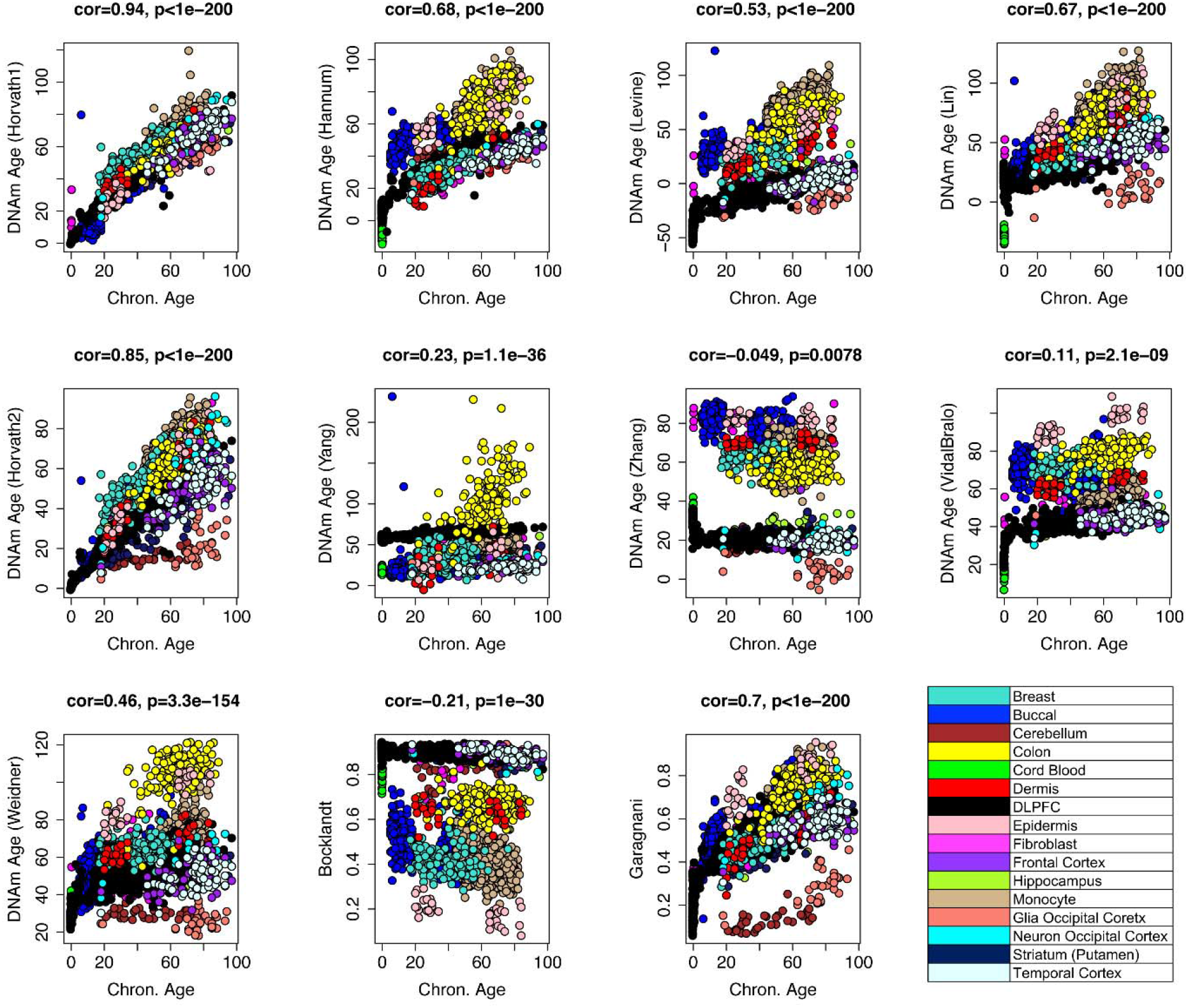
Age correlations for the clock scores across tissue and cell types. DLPFX, dorsolateral prefrontal cortex.

### Transcriptomic signatures of the clocks

To examine the potential functional implications of the various clocks, we examined their transcriptional signals. Fig 5A shows the clustering of the epigenetic clocks based on the log_2_FC values of 5,028 differentially expressed genes (DEGs) that were identified using microarray data from purified monocytes. Results suggested that the age residuals of six clocks—Yang, Hannum, Lin, Levine, Horvath1, and Horvath2—have relatively similar transcriptional signals. Expression associated with these six clocks was further compared to determine the relative strength of their expression signals. As shown in Fig 5B, the log_2_FC for each DEG in association with five clocks (Hannum, Lin, Levine, Yang, and Horvath2) was plotted against the log_2_FC for the association with each DEG and Horvath1. These results suggest that compared to Horvath1, the other five clocks have amplified expression signals (slope >1).

**Fig 5.**
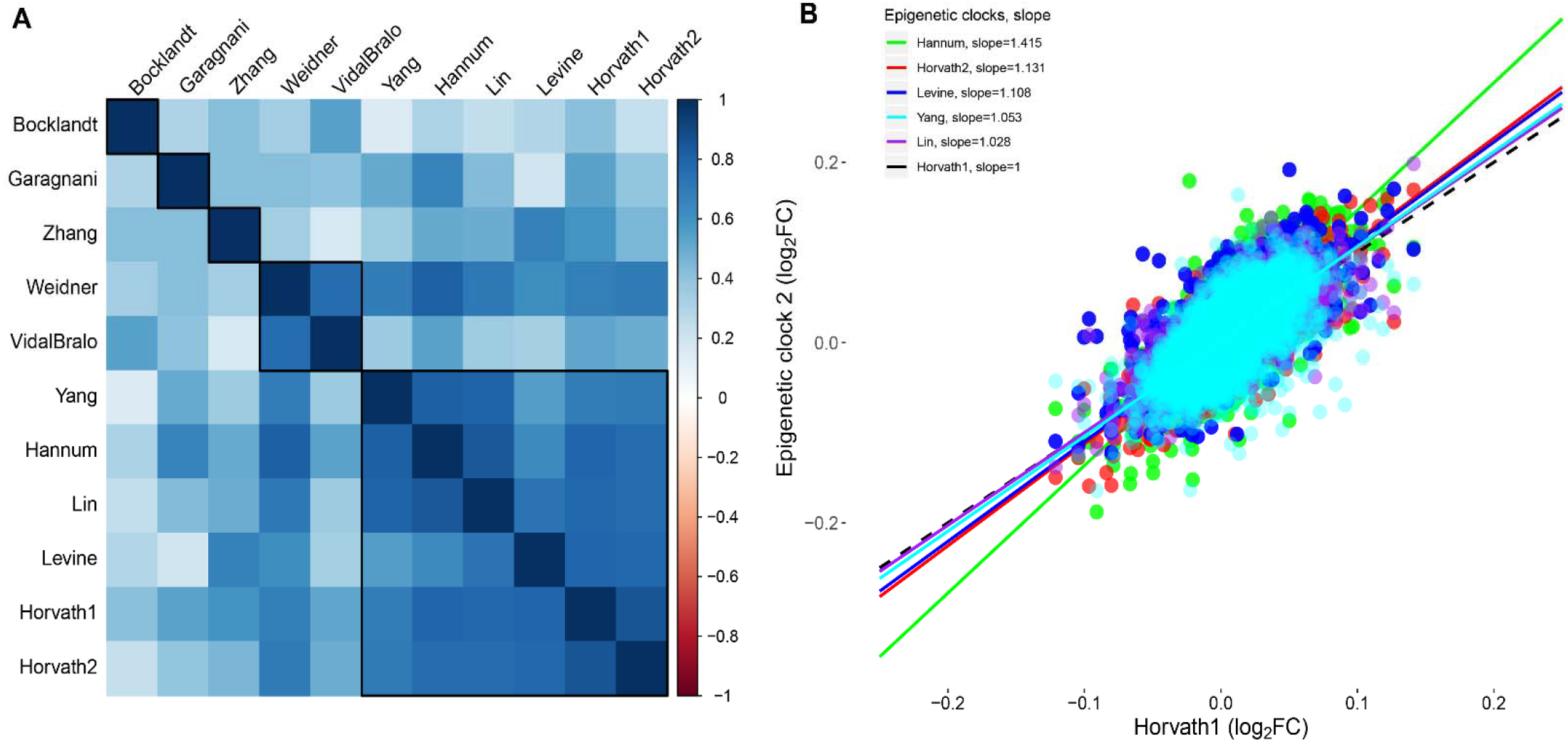
DEGs expression pattern. A. The clustering of the epigenetic clocks based on the log_2_FC values of 5,028 DEGs that were identified out of 10,471 genes in monocytes. B. Comparisons of shared transcriptomic signals for the six clocks (Yang, Hannum, Lin, Levine, Horvath1, and Horvath2) while using the pan-tissue clock by Horvath (Horvath1) as the reference.

Next, we employed weighted gene co-expression network analysis (WGCNA), from which we identified nine co-expression (Fig 6A), ranging in size from 69 for the Magenta module to 838 for the turquoise module. Results based on WGCNA have been shown to be more reproducible and are thought to better capture true biology, compared to traditional differential expression analysis. To test the clock associations with co-expression modules, we estimated eigengene values for each module (equivalent to the first PC among all genes in a given module) and related this value to the clock residuals (Fig 6B, complete results can be found in Fig S3, Additional file 5). Multiple modules displayed robust correlations across various clocks. For example, the blue module is positively associated with the six clocks (age residuals), ranging from β=0.89 (Horvath2) to β=4.14 (Yang). Conversely, the turquoise module negatively relates to age residuals for the six clocks, ranging from β=-0.96 (Horvath2) to β=-3.50 (Yang). The strong correlation (r=-0.86, Additional file 6) between the blue and turquoise modules suggests that they might represent a larger network, but differentially represent overexpressed genes relative to epigenetic age acceleration (blue module), and underexpressed genes relative to epigenetic age acceleration (turquoise module). Overall, most modules showed consistent directions of associations across clocks, with the one exception being the magenta module, which is positively related to Levine and negatively related to Yang.

**Fig 6.**
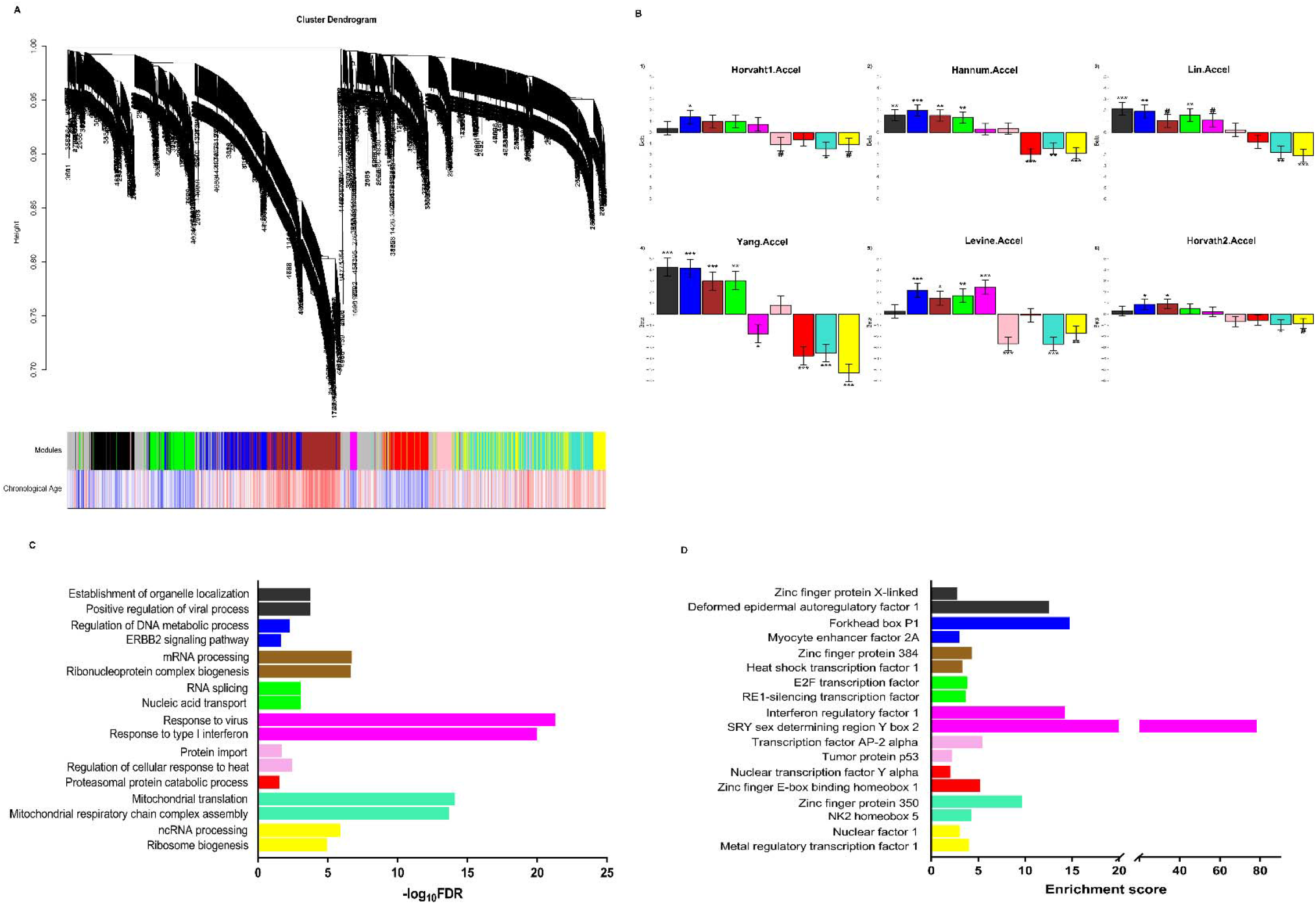
Network analysis of DEGs to unveil transcriptomic signatures of the clocks. A. Network dendrogram from co-expression topological overlap of 5,028 DEGs across the 11 clocks. Color bars show correlation of gene expression with the age acceleration of the clock scores (blue—positive, red—negative). B. Relationship between the eigengene values for each module and the acceleration for the 6 clocks. C. Selected GO enrichment results (complete results are available in Additional file 7 [GO] and 8 [KEGG]). For each module, we selected top two representative biological processes based on q value (FDR), enrichment score, and genes (which should differ for each GO term). D. Selected results from the enrichment analysis for TFBS. Full results from this enrichment analysis can be found in Additional file 11. For each module, we selected top two representative TFs based on P value and enrichment score (Yes/No value>2). If the TF in one module overlapped with that in another module, we selected that with lower P value and higher enrichment score for presentation.

### Co-expression modules that relate to epigenetic aging are enriched for mitochondrial and metabolic pathways

Selected results for GO enrichment analysis are shown in Fig 5D (complete results are available in Additional file 7 [GO] and 8 [KEGG]). Genes in the blue module were enriched for ERBB2 signaling pathway (enrichment score=3.74, FDR=0.024), and regulation of DNA metabolic process (enrichment score=2.65, FDR=0.005). Interestingly, the blue module contained a number of known aging genes, including *IGF1R, SIRT1, NRF2, PIK3R1*, and *ATM*. Genes in the turquoise module were involved in mitochondrial translation, gene expression, respiratory chain complex assembly, and oxidative phosphorylation (OXPHOS). Similar, but slightly weaker enrichments were also observed for genes in the yellow module. Genes in both turquoise and yellow modules exhibited enrichment for NADH dehydrogenase complex assembly. Other potential aging genes in the yellow module included *TFAM, NDUFS3*, and *NDUFS7*. We observed strong connections between the top 20 hub genes within each module (Fig S5 in Additional file 9 and 10), particularly for those in blue and brown modules. Examples of hub genes identified included: *PHF3, PPP1R12A*, and *USP6* in blue module, *IFIT2, STAT1* in magenta module, *NDUFA13* and *TOMM22* in turquoise module.

The enrichment analysis for TFBS suggested critical binding motifs for these modules (Fig 6D, and Additional file 11 and 12). For example, TFs including FOXP1 (Forkhead box P1, enrichment score=14.7) [24, 25] and MEF2 (Myocyte enhancer factor 2A, enrichment score=3.0) [26] were enriched among genes assigned to the blue module, while ZBRK1 (Zinc finger protein 350, enrichment score=9.7) [27], NKX25 (NK2 homeobox 5, enrichment score=4.3) [28], and CREB1 [29] in the turquoise module.

### Mitochondrial Depletion and Epigenetic Aging

Given the enrichment for mitochondrial dysfunction observed in our co-expression modules of interest, we examined the relationship between mitochondrial depletion and epigenetic aging using DNAm data (GSE100249) [30]. We first calculated clock scores for 143B cells with chronically depleted mtDNA (rho0) and 143B controls. Samples included three independent biological replicates for both cases and controls. Clock scores were then standardized in order to be comparable. Analysis of variance revealed that rho0 cells had more than a three- standard deviation increase in epigenetic age for Levine (p=0.006), Lin (p=0.012), and Yang (p=0.013). A slightly weaker increase was also observed for Horvath2 (p=0.044) (Fig S6 in Additional file 13).

## Discussion

In response to the initiative from the National Institute of Health, aimed at identifying valid and reliable biomarkers of aging, a number of potentially promising measures of aging have been developed—perhaps most notably, the epigenetic clocks. Nevertheless, while the growing number of epigenetic clocks have provided some hope that we may be able to successfully track the aging process in vivo using a single measure, there remained a lack of understanding of the characteristics and signals that are either shared between them, or that make each clock distinct. Here, we compared and contrasted the CpG characteristics, age correlations and variances, and transcriptomic signals of 11 unique epigenetic clocks, using data from diverse human tissues and cells. Despite the fact that all the epigenetic clocks were originally developed with the goal of capturing fundamental age-related alterations in the methylation landscape, we found that they each appear somewhat distinct.

The most striking difference is the limited overlap and/or target regions for CpGs included across the clocks. This may not be surprising if in fact hundreds to thousands of CpGs track together with age—representing a single shared biological phenomenon—and each clock randomly selects a small subset of these related CpGs. Another explanation is, given the complexity of the aging process, there are likely distinct types/pathways of epigenetic aging, and that the use of different outcome measures, tissues, and populations in developing the clocks may influence the proportions of each “type” selected by the various clocks. While in general, the proportions of islands, shores, and shelves is comparable among five of the six clocks (Hannum, Lin, Levine, Yang, Horvath1, and Horvath2), the CpGs in these clocks vary in regards to their age correlations in different tissues and cells, suggesting that they may be capturing different biological phenomena. For instance, despite being developed using DNAm data exclusively from whole-blood, CpGs in the clock by Hannum, and to a lesser extent those in the clock by Lin, show very strong age correlations across tissues and/cells (about half positive and half negative). This consistency is not surprising given that it was previously shown that over 70% of age-associated DNA methylation changes are commonly shared across tissue/cell types and in that recent report [31], it was also shown that a high proportion (over 65%) of DNAm changes are associated with age. Conversely, while CpGs in the pan-tissue clock by Horvath (Horvath1) and those in the clock by Levine, show consistency across tissues and cells, many exhibiting little to no age correlation. Given that the Levine clock was trained to predict ‘Phenotypic Age’, which strongly correlates with chronological age, yet at the same time, is meant to differentiate mortality risk among same aged individuals [32], we hypothesize that the large proportion of CpGs in the Levine clock with relatively little to no age correlation, may be capturing stable between-person methylation differences that contribute to differences in the aging process via alterations in biological response or general functioning.

Relatedly, the lack of age correlations for many of the CpGs in Horvath1 may reflect tissue differences. For instance, it has been shown that DNAm can be used to differentiate tissues/cells, and thus when training a multi-tissue predictor, CpGs that differentiate tissues/cells may be included in the score to recalibrate the age prediction so that it best matches chronological age regardless of sample type. While, a goal in developing these measures should always be lack of tissue-specificity, at the same time, different tissue/cells cannot be expected to age at the same rate. Thus, methods that pool tissues may inadvertently be adjusting out some important biological differences. Interestingly, for most clocks we observed that the “rate of aging” appeared slower in brain compared to other tissues. We also did not see the expected increase in variance with age in DLPFC, as was observed in monocytes. Furthermore, as reported by Horvath, we find that the majority of clocks show a slower aging rate in cerebellum, as evidence by a lower slope.

Our results from our transcriptional analysis point to a number of pathways—many already implicated in aging—that may either contribute to epigenetic aging, or be a consequence of it. Further, despite the large aforementioned differences between the clocks, and the fact that they incorporated almost entirely different CpGs, our results suggest that many of these epigenetic clocks may be capturing some shared transcriptional signal/s. We showed that the age-residuals for six epigenetic clocks (Yang, Hannum, Lin, Levine, Horvath1, and Horvath2) had similar associations with gene expression, such that the log2FCs for these various clocks were highly correlated. Further, we also observed relatively consistent associations between clock residuals and co-expression modules. One of the noteworthy co-expression modules was the blue module, which was positively associated with the age residuals for six of the clocks. The blue module was enriched for genes involved in regulation of DNA metabolic processes and epidermal growth factor receptor (EGF) signaling. EGF signaling has been implicated in a number of cancers, particularly breast [33]. In addition to promoting cellular division and differentiation, studies in *C. elegans* suggest a role of EGF in longevity and healthspan [34] and EGF signaling has been shown to have multiple potential points of overlap with the Insulin/IGF-like Signaling (IIS) pathway. Interestingly, studies also suggest EGF receptor (EGFR) may modulate mitochondrial function, through alterations to mitochondrial cytochrome c oxidase subunit II [35, 36].

Another co-expression module of interest was turquoise, which had a strong negative correlation with the blue module, suggesting they were part of the same network, yet distinguished upregulated genes (blue) and downregulated genes (turquoise) in relation to epigenetic aging. The turquoise module was negatively related to seven out of the 11 clocks, and was enriched for nuclear encoded genes involved in mitochondrial transcription, translation, and respiratory chain functions. Age and disease associated downregulation of genes involved in such functions have been reported for a number of organs, including heart [37] and brain [38]. Moreover, the links between mitochondrial function and aging were described more than half a century ago [39] and mitochondrial dysfunction is still considered one of the major hallmarks of aging [2, 5]. However, given how little is known regarding the underlying explanations for DNAm age-alterations, it remains unclear whether changes in DNAm influence mitochondrial functioning, or whether impaired mitochondria contribute to age-related changes in the epigenome. Nevertheless, we observed that Levine DNAmAge—and to some extent Lin, Yang, and Horvath2 DNAmAge—were accelerated in 143B cells with chronically depleted mtDNA (rho0). Such cells, are unable to carry out oxidative phosphorylation, suggesting that the causal pathway may go from mitochondrial dysfunction→epigenetic aging.

Results from our comparative analysis have important implications for future work. First, the majority of epidemiological studies only evaluate Horvath1 and/or Hannum given their notoriety and their growing body of literature. Yet, based on our results showing the differences across existing clocks, it may be more advantageous to either examine multiple clocks simultaneously or to select a clock that best fits the aims of the study. Second, the finding of the amplified transcriptional signal for the Yang clock and the unique gene module represented by others (e.g., Levine) may imply that distinct types of DNAm changes occur in parallel with aging. In moving forward, it will be important to determine whether various types of epigenetic aging have distinct versus common causes. It will also be critical to identify whether each clock has unique downstream functional implications or whether they have additive or multiplicative effects. If the causes and consequences of different types of epigenetic aging are unique, then epigenetic clocks—which consist of a variety of types if DNAm age changes—may exhibit robust age predictions, but at the expense of losing biological specificity and outcome prediction.

In summary, by comparing 11 existing epigenetic clocks, we have observed differences in regards to the CpGs they contain and their characteristics, yet surprisingly consistent transcriptional signals. These results are a first step in uncovering the underlying biology of epigenetic clocks, and in may inform future development of more robust and valid epigenetic biomarkers of aging in both humans and animal models.

## Materials and Methods

### DNA Methylation Data

All data used in this study are publicly available via Gene Expression Omnibus (GEO) and detailed descriptions of these datasets can be found in Additional file 16. Briefly, we used Illumina Infinium 450k DNA methylation data from: breast (GSE101961), buccal (GSE94876), cerebellum (part of GSE89706), colon (GSE101764), dermis (part of GSE51954), dorsolateral prefrontal cortex (DLPFX, GSE74193), epidermis (part of GSE51954), fibroblasts (GSE77135), frontal cortex bulk (part of GSE66351), hippocampus (part of GSE89706), purified monocytes (GSE56046), occipital cortex glia (part of GSE66351), occipital cortex neurons (part of GSE66351), striatum (part of GSE89706), temporal cortex bulk (part of GSE66351), and whole blood (GSE87571).

### Epigenetic clocks

Each of the 11 published clocks considered in this study (Table 1 and Additional file 1), was calculated in accordance with published methods [13, 15, 19, 20, 22, 23, 40-44]. To simplify the description, we used the last name of the first author to refer to each clock. Most of these clocks were developed to predict chronological age in whole blood, with the number of included CpGs ranging from 3 to 513—the exception being the clocks by Bocklandt and Garagnani, which are each based on DNAm for only one CpG. In this study, we also calculated the age acceleration, defined as the residual resulting from a linear model when regressing epigenetic age on chronological age. As mentioned, the age acceleration is meant to reflect between-person and/or between-tissue variably in the rate of aging—whether a person appears older (positive value) or younger (negative value) than expected [11, 12, 43].

### Transcriptomic Data

To better understand the functional signatures of the clocks, we used both transcriptomic (GSE56045) and methylomic (GSE56046) datasets from purified monocytes from 1,202 participants (age 44-83 years) in the Multi-Ethnic Study of Atherosclerosis (MESA) [45] (see details in Additional file 16). For the transcriptomic dataset, in addition to the pre-processing and quality control performed by Reynolds et al (the data provider), we performed other procedures when starting with >47,000 probes: excluded probes with a detection P-value of ≥0.05 (similar to the signal to noise ratio in >50% samples), excluded those targeting putative and/or not well-characterized genes (e.g., gene names starting with FLJ, LOC, and MGC), excluded those with low variance across the samples (<10th percentile), and assigned a mean value to a gene with multiple probes. These procedures resulted in a final dataset containing 10,471 probes with unique Entrez gene IDs that was then used in subsequent analysis.

### Statistical Analyses

The analytic plan is briefly described in Fig 1 and additional details can be found in Additional file 12. In brief, to identify the CpG characteristics (step 1), we: 1) described the overlap in CpG sites and/or CpG blocks across the 11 clocks; 2) compared the CpG types/targets included in each of the clocks (e.g., the proportion of CpG in high density islands, PcG protein targets, DNase I hypersensitive sites); and 3) tested the age correlations and age-specific variance of CpGs within each clock, using data from multiple tissues and cell types, representing the full age range from fetal up to extreme old age.

In step 2, we calculated the 11 epigenetic clocks using a variety of tissues and cell types. We then examine their age correlations across pooled samples, as well as within six tissues/cells— monocytes, DLPFC, colon, fibroblasts, epidermis, glial from occipital cortex, neurons from occipital cortex.

To investigate the functional signatures of these clocks, we related them to transcriptomic data (step 3). Using data from purified monocytes (GSE56046), we first identified genes (denoted as differentially expressed genes, DEGs) that were associated with age residuals for at least one of the 11 epigenetic clocks (FDR<0.05), using R package “limma” [46]. We then compared the gene-specific log_2_FC values corresponding to 11 epigenetic clocks (age residuals), to determine if clocks exhibited shared transcriptomic signals. Using the DEGs, we performed weighted-gene correlation network analysis (WGCNA) [47] to identify co-expression modules. For each module, we estimated the eigengene value—representing the optimal summary of the gene expression profile for gene assigned to that modules—and then related these module eigengenes to the epigenetic clock age residuals. We then performed functional enrichment analysis for Gene Ontology (GO) terms and KEGG pathways for the co-expression modules, using the R package “clusterProfiler” [48]. Next, we identified potential hub genes for each module based on module membership (also referred to as kME or module eigengene based connectivity), defined as the correlation between the module eigengene and the gene expression profile. The top 20 hub genes were then plotted using Cytoscpae (Cytoscape software, version 3.7.0 [49]). Finally, we performed transcription factor binding site (TFBS) enrichment analysis for genes in each module using TRANSFAC^®^ by geneXplain platform (http://genexplain.com/transfac/). Promoter regions (1000 upstream to 100 bp downstream) were interrogated for TF-specific motifs using the TRANSFAC curated TFBS database. By comparing TFBS in genes within each module to the rest of the genome (release 2018.3), we identified overrepresented TFs (P<0.05).

Finally, in step 4, we utilized data from GSE100249 [30] that included DNAm for 143B cells with chronically depleted mtDNA (rho0) ethidium bromide treatment, as well as 143B controls. Samples included three independent biological replicates for both cases and controls. Clock scores were then standardized in order to be comparable, and analysis of variance (ANOVA) was used to compare chronically depleted mtDNA cells to control cells.

All analyses were performed using R version 3.5.1 (2018-07-02).

## References

1. Bauer, U.E., et al., Prevention of chronic disease in the 21st century: elimination of the leading preventable causes of premature death and disability in the USA. Lancet, 2014. 384(9937): p. 45–52.

2. Kennedy, B.K., et al., Geroscience: Linking Aging to Chronic Disease. Cell, 2014. 159(4):p. 709-713.

3. Gladyshev V.N., Aging: progressive decline in fitness due to the rising deleteriome adjusted by genetic, environmental, and stochastic processes. Aging Cell, 2016. 15(4): p. 594–602.

4. Sierra F. and R. Kohanski, Geroscience and the trans-NIH Geroscience Interest Group, GSIG. Geroscience, 2017. 39(1): p. 1–5.

5. Lopez-Otin, C., et al., The hallmarks of aging. Cell, 2013.153(6): p. 1194–217.

6. Florath, I., et al., Cross-sectional and longitudinal changes in DNA methylation with age: an epigenome-wide analysis revealing over 60 novel age-associated CpG sites. (1460-2083 (Electronic)).

7. Johansson, A., U. Enroth S Fau-Gyllensten, and U. Gyllensten, Continuous Aging of the Human DNA Methylome Throughout the Human Lifespan. (1932-6203 (Electronic)).

8. Rakyan, V.K., et al., Human aging-associated DNA hypermethylation occurs preferentially at bivalent chromatin domains. Genome Res, 2010. 20(4): p. 434–9.

9. Teschendorff, A.E., et al., Age-dependent DNA methylotion of genes that are suppressed in stem cells is a hallmark of cancer. Genome research, 2010. 20(4): p. 440–446.

10. Field, A.E., et al., DNA Methylation Clocks in Aging: Categories, Causes, and Consequences. Mol Cell, 2018.71(6): p. 882–895.

11. Horvath S. and K. Raj, DNA methylation-based biomarkers and the epigenetic clock theory of ageing. Nat Rev Genet, 2018.19(6): p. 371–384.

12. Chen, B.H., et al., DNA methylation-based measures of biological age: meta-analysis predicting time to death. Aging (Albany NY), 2016. 8(9): p. 1844–1865.

13. Zhang, Y., et al., DNA methylation signatures in peripheral blood strongly predict all-cause mortality. Nat Commun, 2017. 8:p. 14617.

14. Marioni, R.E., et al., The epigenetic clock is correlated with physical and cognitive fitness in the Lothian Birth Cohort 1936.Int J Epidemiol, 2015. 44.

15. Levine, M.E., et al., An epigenetic biomarker of aging for lifespan and healthspan. Aging (Albany NY), 2018. 10(4): p. 573–591.

16. Levine, M.E., et al., DNA methylation age of blood predicts future onset of lung cancer in the women’s health initiative. Aging (Albany NY), 2015. 7(9): p. 690–700.

17. Ambatipudi, S., et al., DNA methylome analysis identifies accelerated epigenetic ageing associated with postmenopausal breast cancer susceptibility. European Journal of Cancer, 2017. 75: p. 299–307.

18. Zheng, Y., et al., Blood Epigenetic Age may Predict Cancer Incidence and Mortality. EBioMedicine, 2016. 5: p. 68–73.

19. Yang, Z., et al., Correlation of an epigenetic mitotic clock with cancer risk. Genome Biol, 2016. 17(1):p. 205.

20. Horvath, S., et al., Epigenetic clock for skin and blood cells applied to Hutchinson Gilford Progeria Syndrome and ex vivo studies. Aging (Albany NY), 2018. 10(7): p. 1758–1775.

21. Horvath, S., et al., Decreased epigenetic age of PBMCs from Italian semi-supercentenarians and their offspring. Aging (Albany NY), 2015. 7.

22. Weidner, C.I., et al., Aging of blood can be tracked by DNA methylation changes at just three CpG sites. Genome Biology, 2014. 15(2): p. R24.

23. Lin Q. and W. Wagner, Epigenetic Aging Signatures Are Coherently Modified in Cancer. PLoS Genet, 2015. 11(6): p. e1005334.

24. Li, H., et al., FOXP1 controls mesenchymal stem cell commitment and senescence during skeletal aging. J Clin Invest, 2017. 127(4): p. 1241–1253.

25. McClay, J.L., et al., A methylome-wide study of aging using massively parallel seguencing of the methyl-CpG-enriched genomic fraction from blood in over 700 subjects. Hum Mol Genet, 2014. 23(5): p. 1175–85.

26. Chan, S.F., et al., Transcriptional profiling of MEF2-regulated genes in human neural progenitor cells derived from embryonic stem cells. Genom Data, 2015. 3: p. 24–27.

27. Liao, C.C., et al., RB.E2F1 complex mediates DNA damage responses through transcriptional regulation ofZBRK1. J Biol Chem, 2010. 285(43): p. 33134–43.

28. Qu, X.K., et al., A novel NKX2.5 loss-of-function mutation associated with congenital bicuspid aortic valve. Am J Cardiol, 2014. 114(12): p. 1891–5.

29. Arnould, T., et al., CREB activation induced by mitochondrial dysfunction is a new signaling pathway that impairs cell proliferation. EMBO J, 2002. 21(1-2): p. 53–63.

30. Lozoya, O.A., et al., Mitochondrial nicotinamide adenine dinucleotide reduced (NADH) oxidation links the tricarboxylic acid (TCA) cycle with methionine metabolism and nuclear DNA methylation. PLoS Biol, 2018. 16(4): p. e2005707.

31. Zhu, T., et al., Cell and tissue type independent age-associated DNA methylation changes are not rare but common. Aging (Albany NY), 2018. 10(11): p. 3541–3557.

32. Liu, Z., et al., A new aging measure captures morbidity and mortality risk across diverse subpopulations from NHANES IV: A cohort study. PLoS Med, 2018. 15(12): p. e1002718.

33. Normanno, N., et al., Epidermal growth factor receptor (EGFR) signaling in cancer. Gene, 2006. 366(1): p. 2–16.

34. Rongo C., Epidermal growth factor and aging: a signaling molecule reveals a new eye opening function. Aging (Albany NY), 2011. 3(9): p. 896–905.

35. Boerner, J.L., et al., Phosphorylation of Y845 on the epidermal growth factor receptor mediates binding to the mitochondrial protein cytochrome c oxidase subunit II. Mol Cell Biol, 2004. 24(16): p. 7059–71.

36. Demory, M.L., et al., Epidermal growth factor receptor translocation to the mitochondria: regulation and effect. J Biol Chem, 2009. 284(52): p. 36592–604.

37. Preston, C.C., et al., Aging-induced alterations in gene transcripts and functional activity of mitochondrial oxidative phosphorylation complexes in the heart. Mech Ageing Dev, 2008. 129(6): p. 304–12.

38. Mastroeni, D., et al., Nuclear but not mitochondrial-encoded oxidative phosphorylation genes are altered in aging, mild cognitive impairment, and Alzheimer’s disease. Alzheimer’s & dementia: the journal of the Alzheimer’s Association, 2017. 13(5): p. 510–519.

39. Rockstein M. and K.F. Brandt, Enzyme changes in flight muscle correlated with aging and flight ability in the male housefly. Science, 1963. 139(3559): p. 1049–51.

40. Vidal-Bralo, L., Y. Lopez-Golan, and A. Gonzalez, Simplified Assay for Epigenetic Age Estimation in Whole Blood of Adults. Frontiers in genetics, 2016. 7: p. 126–126.

41. Bocklandt, S., et al., Epigenetic predictor of age. PLoS One., 2011. 6.

42. Garagnani, P., et al., Methylation of ELOVL2 gene as a new epigenetic marker of age. Aging Cell, 2012. 11(6): p. 1132–4.

43. Horvath S., DNA methylation age of human tissues and cell types. Genome Biol, 2013. 14(10): p. R115.

44. Hannum, G., et al., Genome-wide methylation profiles reveal quantitative views of human aging rates. Mol Cell, 2013. 49(2): p. 359–367.

45. Reynolds, L.M., et al., Age-related variations in the methylome associated with gene expression in human monocytes and Tcells. Nat Commun, 2014. 5:p. 5366.

46. Ritchie, M.E., et al., limma powers differential expression analyses for RNA-sequencing and microarray studies. Nucleic Acids Res, 2015. 43(7): p. e47.

47. Langfelder P. and S. Horvath, WGCNA: an R package for weighted correlation network analysis. BMC Bioinformatics, 2008. 9: p. 559.

48. Yu, G., et al., clusterProfiler: an R package for comparing biological themes among gene clusters. OMICS, 2012. 16(5): p. 284–7.

49. Shannon, P., et al., Cytoscape: a software environment for integrated models of biomolecular interaction networks. Genome Res, 2003. 13(11): p. 2498–504.

