## Supplementary material for "Comparative analysis of epigenetic aging clocks from CpG characteristics to functional associations": SI materials

**Additional file 1. A brief review of epigenetic aging clocks used in this study**

To date, a few epigenetic clocks have been developed for quantifying human aging (Fig 1 and Table 1). The first one was developed by Bocklandt et al [1] in 2011 in saliva using data from twin pairs. A predictor including two CpGs in the promoter region of EDARADD and NPTX2 explained over 70% of the variance in chronological age, resulting in an average accuracy (average absolute difference from observed chronological age) of about 5 years. In 2012, Garagnani et al [2] reported that one CpG (cg16867657) in ELOVL2 appears to be a promising biomarker of aging (r=0.92) in whole blood using data from 501 persons (9-99 years). One years later, two additional clocks were developed— one by Horvath [3] and one by Hannum et al [4] —which have since become two of the most recognized epigenetic aging clocks in the literature. Both the Horvath and Hannum clocks were developed using penalized regression methods (i.e., elastic net) to train a predictor of chronological age based on DNAm levels at varying numbers of CpGs through the human genome Illumina Infinium arrays. Using about 8,000 samples from 82 datasets (with either 27k or 450k Illumina arrays) that collectively incorporated 51 healthy tissues/cells, Horvath developed a multi-tissue age predictor (often referred to as the pan-tissue clock, including 353 CpGs), which shows age correlations of 0.97 (average accuracy, 2.9) and 0.96 (average accuracy, 3.6) in training and test datasets, respectively[3]. Using 450k array data from 656 persons (19-101 years), Hannum developed an age predictor including 71 CpGs, which shows an age correlation of 0.96 (average accuracy, 3.9) [4].

Later on, other clocks have been proposed using slightly different methods compared to that used by Horvath and Hannum, including 1) quantitative and characteristic-based preselection of CpGs and 2) incorporating other aging outcomes instead of chronological age. Two clocks have been developed by Wolfgang Wagner’s group: the 3 CpG model [5] and the 99 CpG model[6]. Based on 575 pooled blood samples (27k Illumina arrays) from four different studies (0-78 years), 102 CpGs with age correlations over 0.85 were selected first. Weidner et al [5] then selected 3 CpGs based on recursive feature elimination, and conduciveness in a subsequent pyrosequencing analysis, resulting in an age predictor with average accuracy of 5.4 years. This 3 CpGs model was updated for the weights using 450k DNAm array data since it was initially trained on pyrosequencing data [6]. Lin et al [6] validated 99 CpGs out of the 102 preselected age related CpGs in 450k DNAm array, resulting in a 99 CpGs model. Similarly, Vidal-Bralo et al [7] developed an age predictor based on 8 CpG sites, out of a preselected list of the most informative CpGs (with an age correlation over 0.85) using a training set of 390 healthy persons. Yang et al [8] developed a “mitotic clock” using 385 CpGs that meet with three criteria: 1) CpGs should be constitutively unmethylated (or hypomethylated) in any types of fetal tissuesl; 2) CpGs should target the promoters marked by the PRC2 polycomb repressive complex (also known as Polycomb group targets (PCGTs)); 3) CpGs show a trend toward hypermethylation with age. The mitotic clock is estimated as the average DNAm of the 385 CpGs and aims to capture the cellular turnover. Recently, using Illumina 450K and EPIC array data from 10 training datasets, Horvath et al [9] developed a “skin & blood clock” based on 391 CpGs. The 391 CpGs present both on the 450K and EPIC platforms but meet with one of two criteria: 1) having high positive/negative correlation with chronological age in different cell types, or 2) having the least significant correlation with age. This new skin & blood clock was developed for human fibroblasts, keratinocytes, buccal cells, endothelial cells, lymphoblastoid cells, skin, blood, and saliva samples, and can also work for sorted neurons, glia, brain, liver, and even bone samples, outperforming the Horvath pan-tissue clock and Hannum clock.

In contrast, Zhang et al [10] and Levine et al [11] used other aging outcomes other than chronological age to train epigenetic aging measures in their studies. Based on replicated results (58 out of 11,063 CpGs with FDR<0.05) from an epigenome-wide association study (EWAS) for all-cause mortality, Zhang et al further selected 10 CpGs using a LASSO penalized regression method to predict mortality. Two epigenetic aging measures were then proposed—one based on continuous DNAm values of the 10 CpGs, and one based on the sum of aberrant DNAm values (defined as high-risk threshold, either the highest or lowest quartile value) of the 10 CpGs. Levine et al [11] also incorporated mortality prediction, but through a two-step process that initially involved the incorporation of clinical multi-system biomarkers. In step 1, using a Cox proportional elastic net model for mortality, Levine et al selected 9 biomarkers (albumin, creatinine, glucose, [log] C-reactive protein [CRP], lymphocyte percent, mean cell volume, red blood cell distribution width, alkaline phosphatase, and white blood cell count) and therefore developed a novel aging measure (adding chronological age), “phenotypic age” (unit in years), representing the expected age within the population that corresponds to a person’s estimated mortality risk [11, 12]. In step 2, the phenotypic age variable was used as the outcome for training an epigenetic aging clock in whole blood using an elastic net penalized regression approach, resulting in the Levine DNAmPhenoAge including 513 CpGs.

In summary, all these existing clocks are developed by starting with DNAm data for tens to hundreds of thousands of CpG sites across the genome and then training DNAm predictors of age or age-related outcomes using supervised machine learning, resulting in an epigenetic clock including varying numbers of CpGs (typically 50-500).

**Additional file 2.**

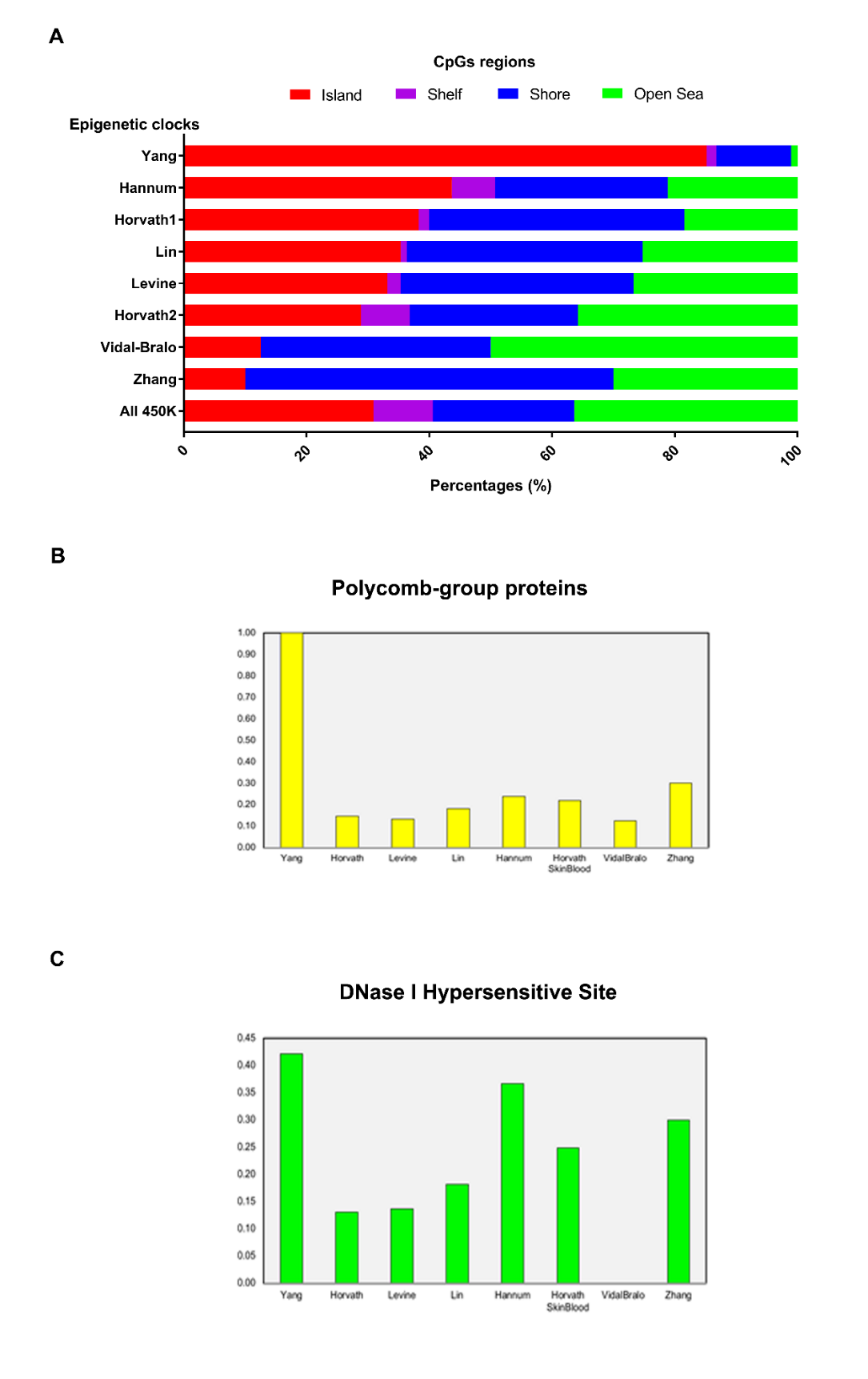

### Fig S1. CpG targets for those included in 11 epigenetic clocks. A. CpGs regions for those included in 11 epigenetic clocks. The results for Bocklandt, Garagnani, and Weidner are not presented. The only one CpG in Bocklandt clock (cg09809672) is located in Shore and the only one CpG in Garagnani clock (cg16867657) in Island. One CpG (cg17861230) in Weidner is located in Island and two (cg02228185 and cg25809905) in Open Sea. B. The proportions of CpGs in polycomb-group (PcG) protein targets. C. The proportions of CpGs in DNase I hypersensitive sites (DHS).

**Additional file 3. Annotations of all CpGs included in 11 epigenetic clocks (separated CSV file)**

**Additional file 4.**

**Fig S2. Age correlations of the clocks in each tissue/cell.**

DLPFX, dorsolateral prefrontal cortex.

From the upper to down, the tissues/cells include monocytes (tan), DLPFC (black), colon (yellow), fibroblasts (magenta), epidermis (pink), glial from occipital cortex (salmon), neurons from occipital cortex (cyan).

**A.**

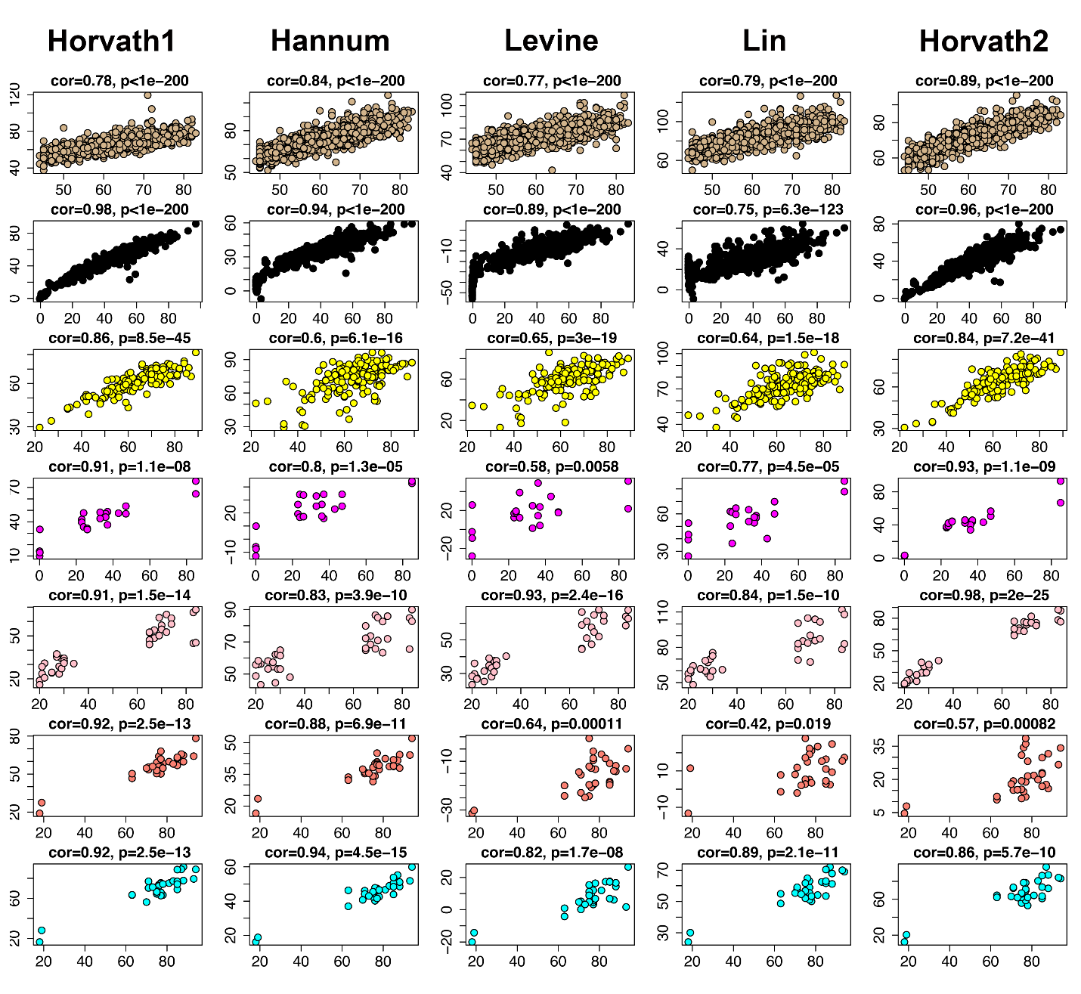

**B.**

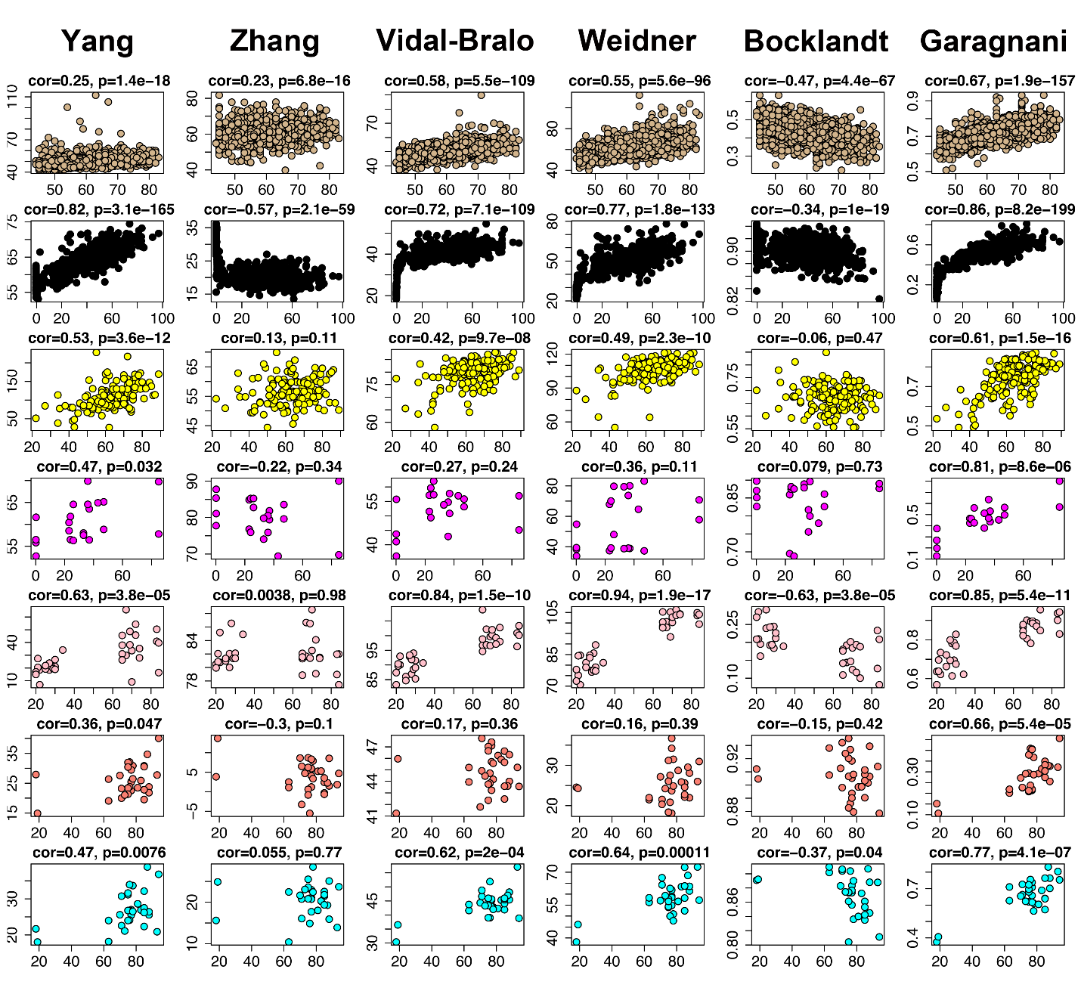

**Additional file 5.**

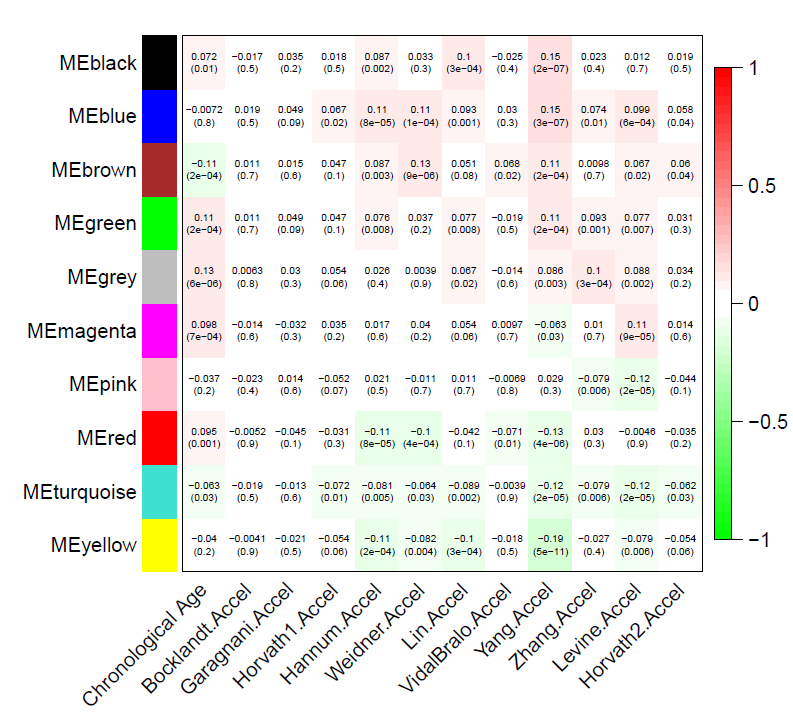

**Fig S3. Module eigengene-traits correlations and p-values.** For each module, an eigengene value were calculated, representing the optimal summary (using principle component analysis in WGCNA) of the gene expression profile. Each cell reports the correlation (and p-value) resulting from correlating module eigengene (rows) to traits (columns, including chronological age and clock accelerations). The table is color-coded by correlation according to the color legend.

**Additional file 6.**

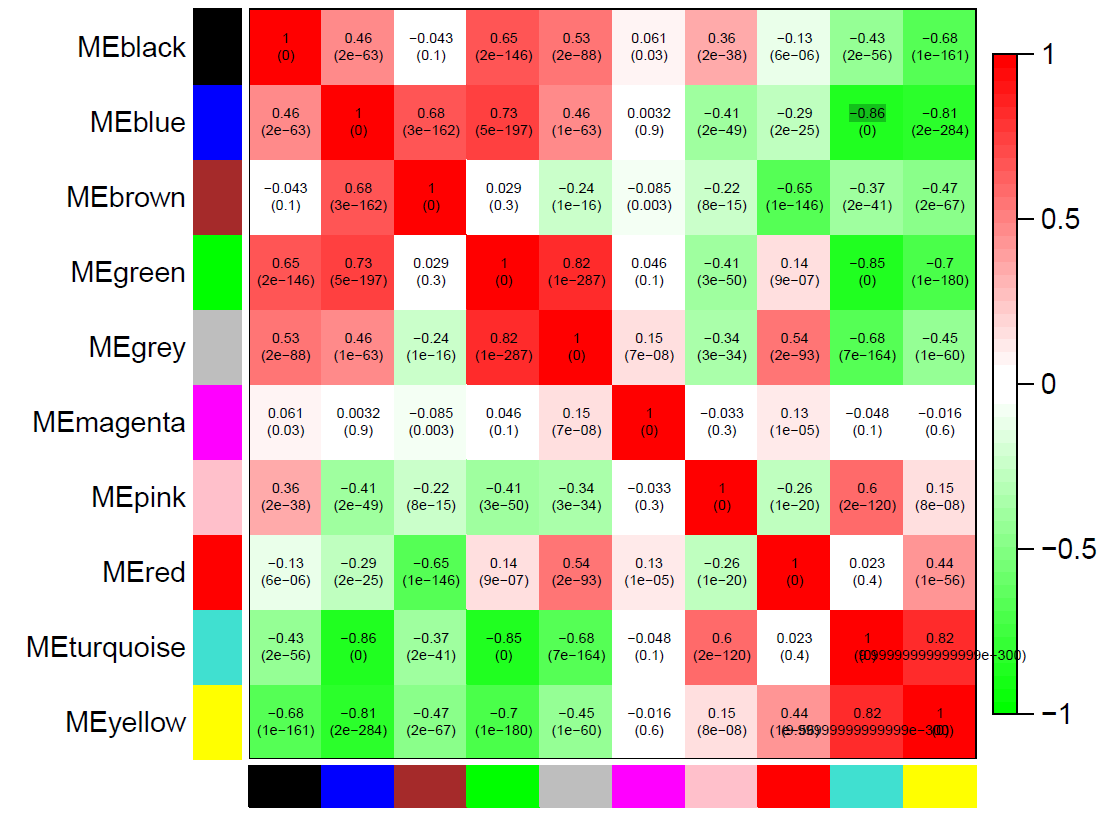

**Fig S4. Module eigengene-module eigengene correlations and p-values.** For each module, an eigengene value were calculated, representing the optimal summary (using principle component analysis in WGCNA) of the gene expression profile. Each cell reports the correlation (and p-value) resulting from correlating one module eigengene (rows) to another module eigengene (columns). The table is color-coded by correlation according to the color legend.

**Additional file 7.**

Results for the GO enrichment analysis for genes in these modules (separated Excel file)

**Additional file 8.**

Results for the KEGG pathway analysis for genes in these modules (separated Excel file)

**Additional file 9.** kME for all genes in each module (separated CSV file)

kME is a quantitative measure (also referred to as module membership, or module eigengene based connectivity), defined as the correlation between the module eigengene and the gene expression profile.

**Additional file 10.**

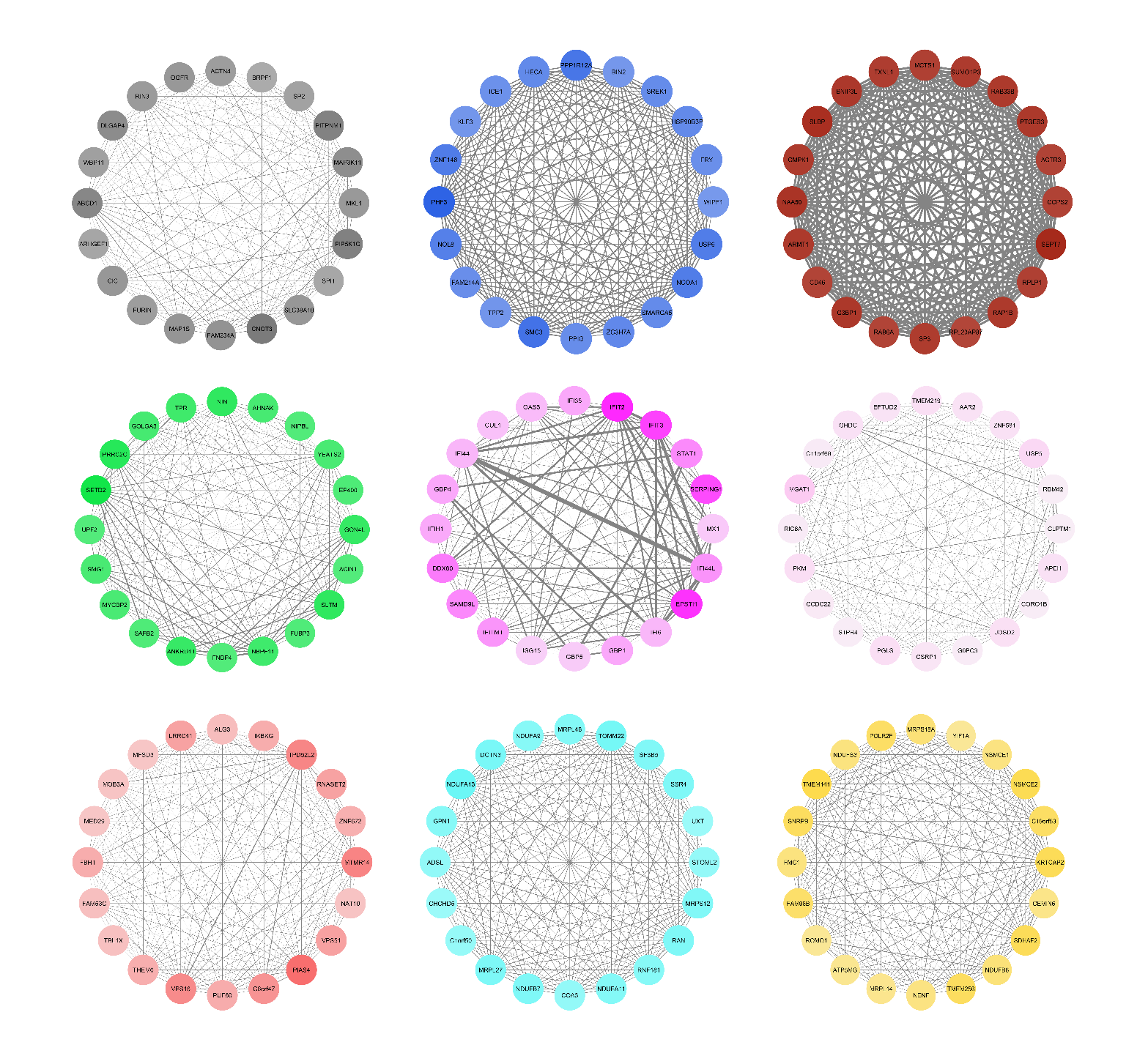

**Fig S5. The top 20 hub genes within each module.**

Full results for kME for all genes in each module can be found in Additional file 9. Edges are weighted by the strength of correlation between genes.

**Additional file 11.**

Detailed results for the TFBS enrichment analysis (separated Excel file)

**Additional file 12.**

**More details on data analysis and results**

**Step 3**

For WGCNA, we designated a “signed” network, and employed a thresholding power of 11 (picked using pickSoftThreshold function) and dynamic tree cut. For each resultant module, WGCNA used using principle component analysis as default to estimate the eigengene value—representing the optimal summary of the gene expression profile for that modules.

**Functional annotation analysis**

For functional annotation analysis (enrichment analysis for Gene Ontology (GO) terms and KEGG pathways), we mainly used the R package “clusterProfiler” [13] as mentioned in the text. However, we also used WebGestalt (<http://www.webgestalt.org/option.php>, a web-based pathway analysis tool) [14] to validate our results. We used the default minimum pathway size (n=5) and significance level of FDR <0.05.

**Additional analysis**

To capture more age related genes in analysis for transcriptomic signatures of the clocks, we performed an additional analysis, in which we included DEGs that were differentially expressed in association with either at least one of the original epigenetic clocks or with the age residuals. We then repeated the step 3 for this group of DEGs. The results did not change substantially (Additional file 14 and 15) although more numbers of modules were identified. For instance, the turquoise module was similar to the turquoise module in the main analysis, exhibiting strong correlations with Yang and Levine clock. The pink module was similar to the magenta module in the main analysis, only showing strong negative correlations with the Levine clock.

**Results for the enrichment analysis for TFBS**

Here we described more interesting results from the enrichment analysis for TFBS (Additional file 9). In addition to the blue module, we found that TFs including E2F (E2F transcription factor, enrichment score=3.8) [15] and REST (RE1-silencing transcription factor, enrichment score=3.7) [16] were enriched among genes assigned to the green module; IRF1 (Interferon regulatory factor 1, enrichment score=14.2) [17, 18] and SOX2 (SRY sex determining region Y box 2, enrichment score=78.3) [19, 20] for the magenta module; and NFY (Nuclear transcription factor Y alpha, enrichment score=2.0) [21] and DELTAEF1 (Zinc finger E-box binding homeobox 1, enrichment score=5.2) [22] for the red module. Notable that CREB1 that involves in mitochondrial dysfunction and cell proliferation [23] was also found in our turquoise module with the smallest P value despite with enrichment score of 1.5. Similar observations were found for RNF96/TRIM28 [24] in blue module, CPBP/KLF6 [25] and EGR1 [26] in green module, and ARID3A/DRIL1 [27] and STAT1 [28] in magenta module.

**Additional file 13.**

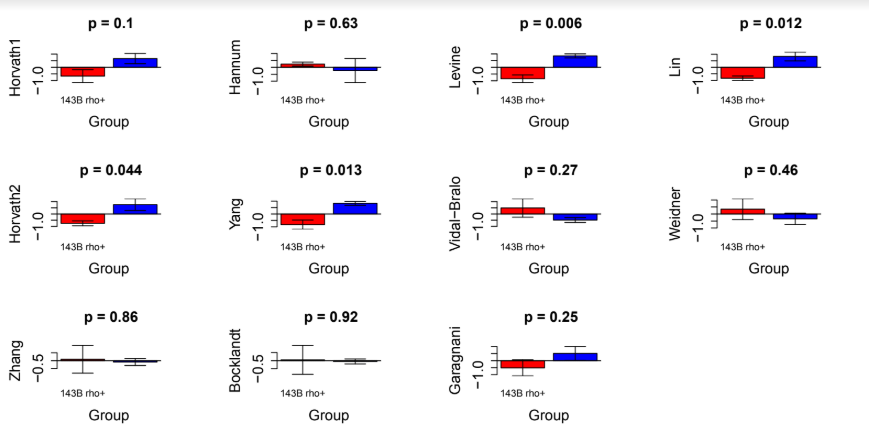

**Fig S6. Mitochondrial Depletion and Epigenetic Aging**

**Additional file 14.**

**Fig S3. Dendrogram for 7,158 DEGs** (separated PDF file)

These 7,158 DEGs that were differentially expressed in association with either at least one of the original epigenetic clocks or one acceleration of the 11 epigenetic clocks (i.e., after adjusting for chronological age). We used similar parameters as our main analysis for 5028 DEGs to draw this dendrogram. A total of 14 gene modules (except the grey) were identified. For color bar below shows the relationship of these genes with epigenetic clocks and clocks accelerations, the closer to “blue”, the higher positive relations, the closer to “red”, the high negative relations.

**Additional file 15.**

**Fig S4. Module eigengene-traits correlations and p-values based on 7,158 DEGs** (separated PDF file)

These 7,158 DEGs that were differentially expressed in association with either at least one of the original epigenetic clocks or one acceleration of the 11 epigenetic clocks (i.e., after adjusting for chronological age). For each module, an eigengene value were calculated, representing the optimal summary (using principle component analysis in WGCNA) of the gene expression profile. Each cell reports the correlation (and p-value) resulting from correlating module eigengene (rows) to traits (columns, including chronological age, clock scores, and clock accelerations). The table is color-coded by correlation according to the color legend.

**Additional file 16. Datasets used in this study**

**Datasets for DNA methylation**

**Breast (GSE101961)**

This dataset measured DNA methylation in the normal breast tissues using the Illumina Infinium 450K Human Methylation Beadchip. It consists of 121 cancer-free women (mean age: 38 years, range: 17-76 years), provided by Song et al [29].

**Buccal (GSE94876)**

This dataset measured DNA methylation in the buccal cells using the Illumina Infinium 450K Human Methylation Beadchip. It includes 120 generally healthy adult males (mean age: 38 years, range: 17-76 years) from the smokers (SMK), moist snuff consumers (MSC), and Non-Tobacco Consumer (NTC) cohorts (40 participants/cohort). It is provided by Jessen et al [30].

**Cerebellum (part of GSE89706)**

This dataset measured DNA methylation in the post-mortem brain samples, representing tissue from four brain regions (prefrontal cortex, striatum, hippocampus and cerebellum), using the Illumina Infinium 450K Human Methylation Beadchip. It includes 262 samples from 41 schizophrenia patients and 47 controls. Here we used the control cerebellum subsamples (n=33, mean age: 45.12 years, range: 21-72 years). This dataset is provided by Viana et al [31].

**Colon (GSE101764)**

This dataset measured DNA methylation in 149 mucosa and 112 colorectal cancer tissues using the Illumina Infinium 450K Human Methylation Beadchip. In this study, we used the 149 mucosa colorectal mucosa samples (mean age: 63 years, range: 22-89 years). This dataset was provided by Barrow et al [32].

**Dermis (part of GSE51954)**

This dataset measured DNA methylation in the dermal and epidermal samples using the Illumina Infinium 450K Human Methylation Beadchip. It consists of 40 dermal and 38 epidermal samples. Here we used the dermis subsample (mean age: 50 years, range: 20-90 years). This dataset is provided by Vandiver et al [33].

**Dorsolateral prefrontal cortex (DLPFX, GSE74193)**

This dataset measured DNA in the dorsolateral prefrontal cortex brain tissue using the Illumina Infinium 450K Human Methylation Beadchip. It consists of 675 samples (mean age: 35 years, range: -0.5-97 years), provided by Jaffe et al [34].

**Epidermis (part of GSE51954)**

This dataset has been described above. Here we used the 38 epidermal samples (mean age: 51 years, range: 20-90 years).

**Fibroblasts (GSE77135)**

This dataset measured in postmortem dural and scalp fibroblasts using the Illumina Infinium 450K Human Methylation Beadchip. it consists of 21 samples (11 intrinsically matched pairs of dural and scalp fibroblasts, 1 was removed in preprocessing) (mean age: 31.7 years, range: 0-85 years), and is provided by Ivanov et al [35].

**Frontal Cortex bulk (part of GSE66351)**

This dataset measured in postmortem human brains using the Illumina Infinium 450K Human Methylation Beadchip. It consists of 63 samples from bulk cells in the frontal cortex (mean age: 73.9 years, range: 18-97 years). This dataset is provided by Gasparoni et al [36].

**Hippocampus (part of GSE89706)**

This dataset has been described above. Here we used the control Hippocampus subsamples (n=27, mean age: 62 years, range: 25-95 years).

**Monocytes (GSE56046)**

This dataset measured DNA methylation in purified monocytes using the Illumina Infinium 450K Human Methylation Beadchip. These samples were from 1,202 participants from the Multi-Ethnic Study of Atherosclerosis (MESA) study ^[37]^. As both the methylation and transcriptomic (GSE56045) datasets from purified monocytes from these participants were used in this study, we briefly described the MESA study. The MESA is a population cohort study with the aim of examining the prevalence, correlates, and progression of subclinical cardiovascular disease since 2000. Data on socio-demographic, lifestyles, nutrition, laboratory, extensive clinical, and medication were collected by five clinic visits. The sample used for transcriptome and methylation analysis were from April 2010 to February 2012 examination (Exam 5) of 1,264 randomly selected participants from four MESA field centers (Baltimore, MD; Forsyth County, NC; New York, NY; and St. Paul,MN) [37]. The details on blood specimen collections, purification of CD14+ monocytes, DNA/RNA extraction, global expression quantification, epigenome-wide methylation quantification, quality control and pre-processing of microarray data were provided elsewhere [37]. Regarding participants included in the methylation and transcriptomic datasets, their mean age was 60 years, with the range of 44 to 83 years. This dataset also includes covariates such as race, gender, study site, and residual sample contamination (i.e., separate enrichment scores for neutrophils, B cells, T cells, and natural killer cells) for further monocyte data analysis. This dataset was provided by Reynolds et al ^[37]^.

**Occipital Cortex Glia and Neurons (part of GSE66351)**

This dataset has been described above. Here we used the sorted cells (glia and neurons) from the occipital cortex (n=31 each cell type, mean age: 74.77 years, range: 18-94 years).

**Striatum (part of GSE89706)**

This dataset has been described above. Here we used the control Striatum (putamen) subsamples (n=82, mean age: years, range: years).

**Temporal cortex bulk (part of GSE66351)**

This dataset has been described above. Here we used the Temporal cortex bulk subsamples (n=?, mean age: 56 years, range: 21-96 years).

**Datasets for transcriptome**

**Monocytes (GSE56045)**

This dataset measured transcriptomic profile in purified monocytes using the Illumina HumanHT-12 v4 Expression BeadChip. The samples were from the same participants in the dataset (GSE56046) above. This dataset was provided by Reynolds et al [37]^,^[38].

**References**

1.	Bocklandt, S., et al., *Epigenetic predictor of age.* PLoS One., 2011. **6**.

2.	Garagnani, P., et al., *Methylation of ELOVL2 gene as a new epigenetic marker of age.* Aging Cell, 2012. **11**(6): p. 1132-4.

3.	Horvath, S., *DNA methylation age of human tissues and cell types.* Genome Biol, 2013. **14**(10): p. R115.

4.	Hannum, G., et al., *Genome-wide methylation profiles reveal quantitative views of human aging rates.* Mol Cell, 2013. **49**(2): p. 359-367.

5.	Weidner, C.I., et al., *Aging of blood can be tracked by DNA methylation changes at just three CpG sites.* Genome Biology, 2014. **15**(2): p. R24.

6.	Lin, Q. and W. Wagner, *Epigenetic Aging Signatures Are Coherently Modified in Cancer.* PLoS Genet, 2015. **11**(6): p. e1005334.

7.	Vidal-Bralo, L., Y. Lopez-Golan, and A. Gonzalez, *Simplified Assay for Epigenetic Age Estimation in Whole Blood of Adults.* Frontiers in genetics, 2016. **7**: p. 126-126.

8.	Yang, Z., et al., *Correlation of an epigenetic mitotic clock with cancer risk.* Genome Biol, 2016. **17**(1): p. 205.

9.	Horvath, S., et al., *Epigenetic clock for skin and blood cells applied to Hutchinson Gilford Progeria Syndrome and ex vivo studies.* Aging (Albany NY), 2018. **10**(7): p. 1758-1775.

10.	Zhang, Y., et al., *DNA methylation signatures in peripheral blood strongly predict all-cause mortality.* Nat Commun, 2017. **8**: p. 14617.

11.	Levine, M.E., et al., *An epigenetic biomarker of aging for lifespan and healthspan.* Aging (Albany NY), 2018. **10**(4): p. 573-591.

12.	Liu, Z., et al., *Phenotypic Age: a novel signature of mortality and morbidity risk.* bioRxiv, 2018.

13.	Yu, G., et al., *clusterProfiler: an R package for comparing biological themes among gene clusters.* OMICS, 2012. **16**(5): p. 284-7.

14.	Wang, J., et al., *WebGestalt 2017: a more comprehensive, powerful, flexible and interactive gene set enrichment analysis toolkit.* Nucleic Acids Res, 2017. **45**(W1): p. W130-W137.

15.	Iaquinta, P.J. and J.A. Lees, *Life and death decisions by the E2F transcription factors.* Curr Opin Cell Biol, 2007. **19**(6): p. 649-57.

16.	Hu, Y., et al., *RE1 silencing transcription factor (REST) negatively regulates ALL1-fused from chromosome 1q (AF1q) gene transcription.* BMC Mol Biol, 2015. **16**: p. 15.

17.	Yang, H., et al., *Histone deacetylase sirtuin 1 deacetylates IRF1 protein and programs dendritic cells to control Th17 protein differentiation during autoimmune inflammation.* J Biol Chem, 2013. **288**(52): p. 37256-66.

18.	Langlais, D., L.B. Barreiro, and P. Gros, *The macrophage IRF8/IRF1 regulome is required for protection against infections and is associated with chronic inflammation.* J Exp Med, 2016. **213**(4): p. 585-603.

19.	Amador-Arjona, A., et al., *SOX2 primes the epigenetic landscape in neural precursors enabling proper gene activation during hippocampal neurogenesis.* Proc Natl Acad Sci U S A, 2015. **112**(15): p. E1936-45.

20.	Ocampo, A., et al., *In Vivo Amelioration of Age-Associated Hallmarks by Partial Reprogramming.* Cell, 2016. **167**(7): p. 1719-1733 e12.

21.	Romier, C., et al., *The NF-YB/NF-YC structure gives insight into DNA binding and transcription regulation by CCAAT factor NF-Y.* J Biol Chem, 2003. **278**(2): p. 1336-45.

22.	Jiang, Y., et al., *Zinc finger E-box-binding homeobox 1 (ZEB1) is required for neural differentiation of human embryonic stem cells.* J Biol Chem, 2018. **293**(50): p. 19317-19329.

23.	Arnould, T., et al., *CREB activation induced by mitochondrial dysfunction is a new signaling pathway that impairs cell proliferation.* EMBO J, 2002. **21**(1-2): p. 53-63.

24.	Santos, J. and J. Gil, *TRIM28/KAP1 regulates senescence.* Immunol Lett, 2014. **162**(1 Pt B): p. 281-9.

25.	Hsieh, P.N., et al., *Aging and the Kruppel-like factors.* Trends Cell Mol Biol, 2017. **12**: p. 1-15.

26.	Krones-Herzig, A., E. Adamson, and D. Mercola, *Early growth response 1 protein, an upstream gatekeeper of the p53 tumor suppressor, controls replicative senescence.* Proc Natl Acad Sci U S A, 2003. **100**(6): p. 3233-8.

27.	Rhee, C., et al., *Arid3a is essential to execution of the first cell fate decision via direct embryonic and extraembryonic transcriptional regulation.* Genes Dev, 2014. **28**(20): p. 2219-32.

28.	Bancerek, J., et al., *CDK8 kinase phosphorylates transcription factor STAT1 to selectively regulate the interferon response.* Immunity, 2013. **38**(2): p. 250-62.

29.	Song, M.A., et al., *Landscape of genome-wide age-related DNA methylation in breast tissue.* Oncotarget, 2017. **8**(70): p. 114648-114662.

30.	Jessen, W.J., M.F. Borgerding, and G.L. Prasad, *Global methylation profiles in buccal cells of long-term smokers and moist snuff consumers.* Biomarkers, 2018. **23**(7): p. 625-639.

31.	Viana, J., et al., *Schizophrenia-associated methylomic variation: molecular signatures of disease and polygenic risk burden across multiple brain regions.* Hum Mol Genet, 2017. **26**(1): p. 210-225.

32.	Barrow, T.M., et al., *Smoking is associated with hypermethylation of the APC 1A promoter in colorectal cancer: the ColoCare Study.* J Pathol, 2017. **243**(3): p. 366-375.

33.	Vandiver, A.R., et al., *Age and sun exposure-related widespread genomic blocks of hypomethylation in nonmalignant skin.* Genome Biol, 2015. **16**: p. 80.

34.	Jaffe, A.E., et al., *Mapping DNA methylation across development, genotype and schizophrenia in the human frontal cortex.* Nat Neurosci, 2016. **19**(1): p. 40-7.

35.	Ivanov, N.A., et al., *Strong Components of Epigenetic Memory in Cultured Human Fibroblasts Related to Site of Origin and Donor Age.* PLoS Genet, 2016. **12**(2): p. e1005819.

36.	Gasparoni, G., et al., *DNA methylation analysis on purified neurons and glia dissects age and Alzheimer's disease-specific changes in the human cortex.* Epigenetics Chromatin, 2018. **11**(1): p. 41.

37.	Reynolds, L.M., et al., *Transcriptomic profiles of aging in purified human immune cells.* BMC Genomics, 2015. **16**: p. 333.

38.	Reynolds, L.M., et al., *Age-related variations in the methylome associated with gene expression in human monocytes and T cells.* Nat Commun, 2014. **5**: p. 5366.
